# Development of a honey bee RNA virus vector based on the genome of Deformed wing virus

**DOI:** 10.1101/2020.02.18.954958

**Authors:** Eugene V. Ryabov, Krisztina Christmon, Matthew C. Heerman, Francisco Posada-Florez, Robert L. Harrison, Yanping Chen, Jay D. Evans

## Abstract

We developed a honey bee RNA-virus vector based on the genome of a picorna-like Deformed wing virus (DWV), the main viral pathogen of the honey bee (*Apis mellifera*). To test the potential of DWV to be utilized as a vector, the 717 nt sequence coding for the enhanced green fluorescent protein (eGFP), flanked by the peptides targeted by viral protease, was inserted into an infectious cDNA clone of DWV in-frame between the leader protein and the virus structural protein VP2 genes. The *in vitro* RNA transcripts from *egfp*-tagged DWV cDNA clones were infectious when injected into honey bee pupae. Stable DWV particles containing genomic RNA of the recovered DWV with *egfp* inserts were produced, as evidenced by cesium chloride density gradient centrifugation. These particles were infectious to honey bee pupae when injected intra-abdominally. Fluorescent microscopy showed GFP expression in the infected cells and Western blot analysis demonstrated accumulation of free eGFP rather than its fusions with DWV LP and/or VP2 proteins. Analysis of the progeny *egfp*-tagged DWV showed gradual accumulation of genome deletions for *egfp*, providing estimates for the rate of loss of a non-essential gene an insect RNA virus genome during natural infection.

## 1. Introduction

Vectors based on the genomes of RNA viruses of animals [1,2] or plants [3,4] have been widely used in research and in biotechnology since the 1990s. RNA vectors provide transient peptide expression at high levels for research and for industrial protein production [4] without the generation of transgenic hosts. Moreover, RNA virus vectors allow the expression of proteins which could be toxic or affect host development, a complication for transgenic approaches. Another unique feature of RNA virus vectors is their use for the induction of RNA interference (RNAi) due to the generation of double-stranded (ds) RNA replication intermediates [5]. Importantly, virus-based vectors that express green fluorescent protein (GFP) or other reporters enable profound studies of viruses and their interactions with hosts [6].

Insect-specific RNA virus vectors have been designed using genomes of the arboviruses, members of Flavivirus, Alphavirus and Nodavirus groups [1,7,8]. Currently, there are no RNA vectors based on genomes of viruses naturally infecting honey bees. While GFP-tagged vector based on a cloned Sindbis virus, a member of the Alphavirus group, was successfully used in honey bees [9], development of reporters tied to an infective honey bee virus would provide a novel system to investigate virus and host interactions.

In this study, we develop a virus expression system for the widespread virus DWV type A [10] using a recently designed infectious cDNA clone of genomic RNA of this virus [11]. We identified a site in the DWV genomic RNA insertion into which of an enhanced green fluorescent protein (eGFP)-coding sequence [12], corresponding to 7 % of the DWV RNA genome, did not affect virus replication. We demonstrated encapsidation of DWV RNA following *egfp* gene insertion, and showed expression of the *egfp* reporter in the cells of honey bee pupae infected with the *egfp*-tagged DWV RNA vectors by using fluorescent microscopy and Western blot. Analysis of genetic stability of the DWV vector provided novel insights into the ability of RNA viruses to retain non-essential sequences, contributing to a better understanding of their evolution.

## 2. Materials and Methods

### 2.1. Construction of the egfp-tagged DWV cDNA clones and generation of the clone-derived viruses

The full-length infectious cDNA clone of DWV type A, pDWV-304 (GenBank accession number MG831200) [11] was used to design eGFP-expressing constructs, pDWV-L-GFP and pDWV-S-GFP (Figure 1; Text S1). Cloning included generation of sections of DWV cDNA with *egfp* insertions at the LP-VP2 border by overlap extension PCR and subcloning, using pDWV-304 [11] and pAcP(+)IE1-eGFP [13] as the templates for DWV and *egfp* amplification, respectively, and the primers listed in Table S1. Cloning procedure is described in detail in Text S1.

**Figure 1.**
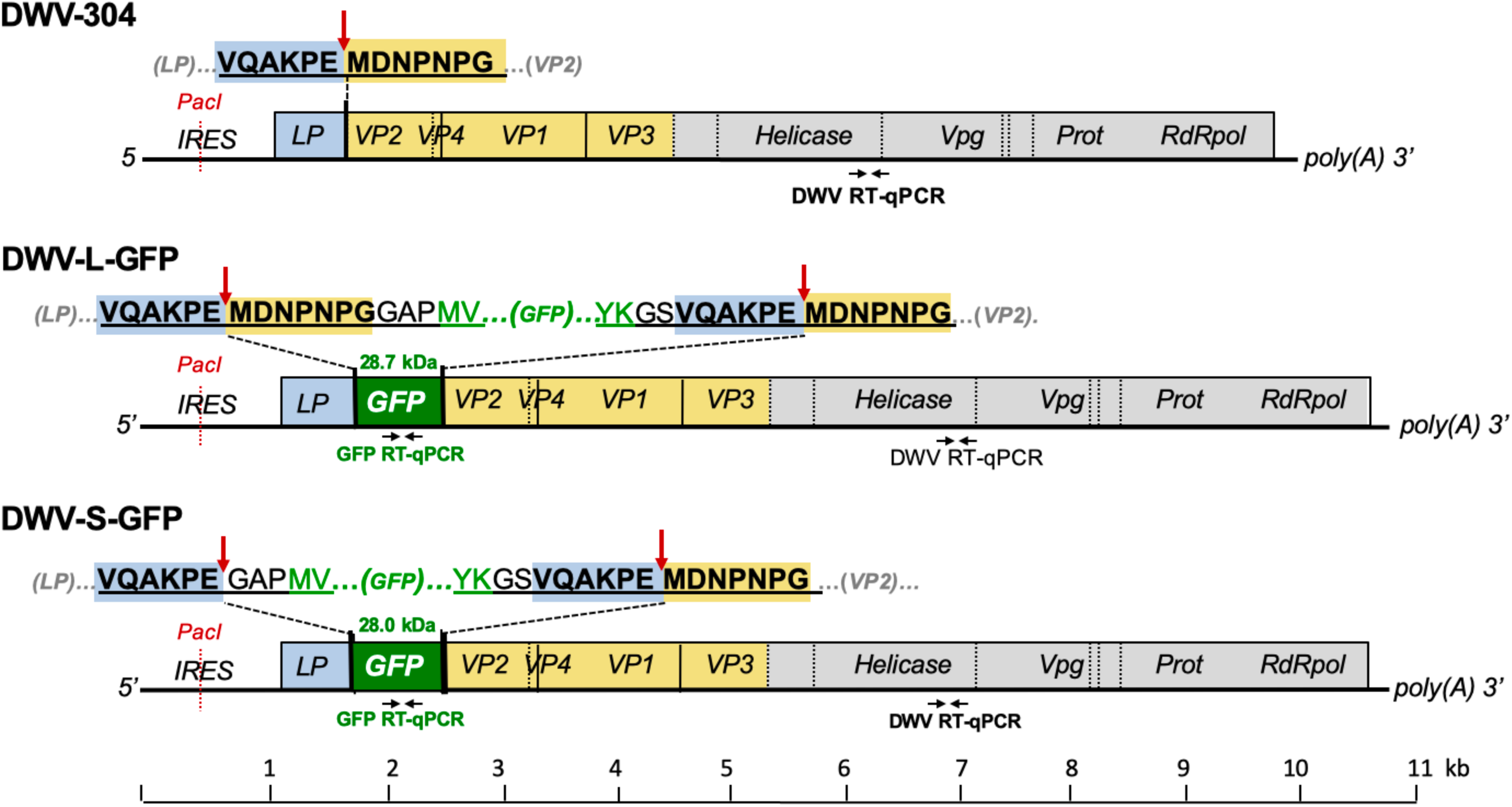
Design of the GFP-tagged DWV cDNA clones. Sequences of the peptide linking LP-VP2 and the GFP-flanking peptides are shown above the genetic maps. Arrows indicate the tentative cleavage peptides targeted by the DWV 3C protease, underscored are DWV-derived peptides. Positions of the DWV- and GFP-specific RT-qPCR primers are indicated. DWV, deformed wing virus; LP, leader protein; GFP, green fluorescent protein; Prot, viral 3C protease; RdRpol, RNA-dependent RNA polymerase; RT-PCR, reverse-transcription PCR; VP, structural viral protein; Vpg, genome-linked protein.

The *egfp*-tagged DWV cDNA plasmid constructs were linearized using the *Pme*I restriction site located at the 3′ end of DWV cDNA, downstream of the poly A sequence, to produce the templates for a full-length 10.9-kb transcripts. The *in vitro* RNA transcripts from pDWV-L-GFP, pDWV-S-GFP and the parental pDWV-304 were produced using HiScribe T7 High Yield RNA Synthesis Kit (New England Biolabs) according to the manufacturer’s instructions. After 3 hours of incubation at +37 °C, the DNA templates were digested using TURBO DNase (Life Technologies). The *in vitro* RNA transcripts purified using the RNeasy mini kit (Qiagen), 13.2 × 10^12^ copies in 10 µL of PBS, or 10 µL of PBS were injected intra-abdominally into the hemolymph of honey bee pupae at pink eye stage which were not exposed to *Varroa* mite feeding using syringes with a 0.3-mm needle G31 (BD Micro-Fine) as describer previously [11]. The extracts containing DWV-L-GFP and DWV-S-GFP virus particles were produced by homogenizing individual transcript-injected pupae with 1.5 mL of PBS at 72 hpi and subjected to three cycles of freeze-thawing, preparation included clarification by centrifugation at 3,000 g for 5 minutes and filtration through a 0.22 µm nylon filter (Millipore). The DWV concentration in the extracts was quantified by RT-qPCR, and the extracts were stored at −80 °C prior to use.

### 2.2. Honey bees

Honey bee pupae were obtained from strong colonies with low (less than 1 %) *Varroa* mite infestation rate from the apiaries located in the USDA Bee Research Laboratory (BRL) in Beltsville, Maryland (sourced in September 2019), and in the USDA Honey Bee Breeding, Genetics and Physiology Research Laboratory in Baton Rouge, Louisiana USA (sourced in November 2019). Pupa at the pink-eyed stage were pulled out of cells using soft tweezers 3 hours prior their use in the injection experiments which were carried out as described previously [11]. Newly emerged honey bees with deformed wings sourced in Maryland in 2019 were used as source of wild-type DWV for Fig. 7B.

**Figure 2.**
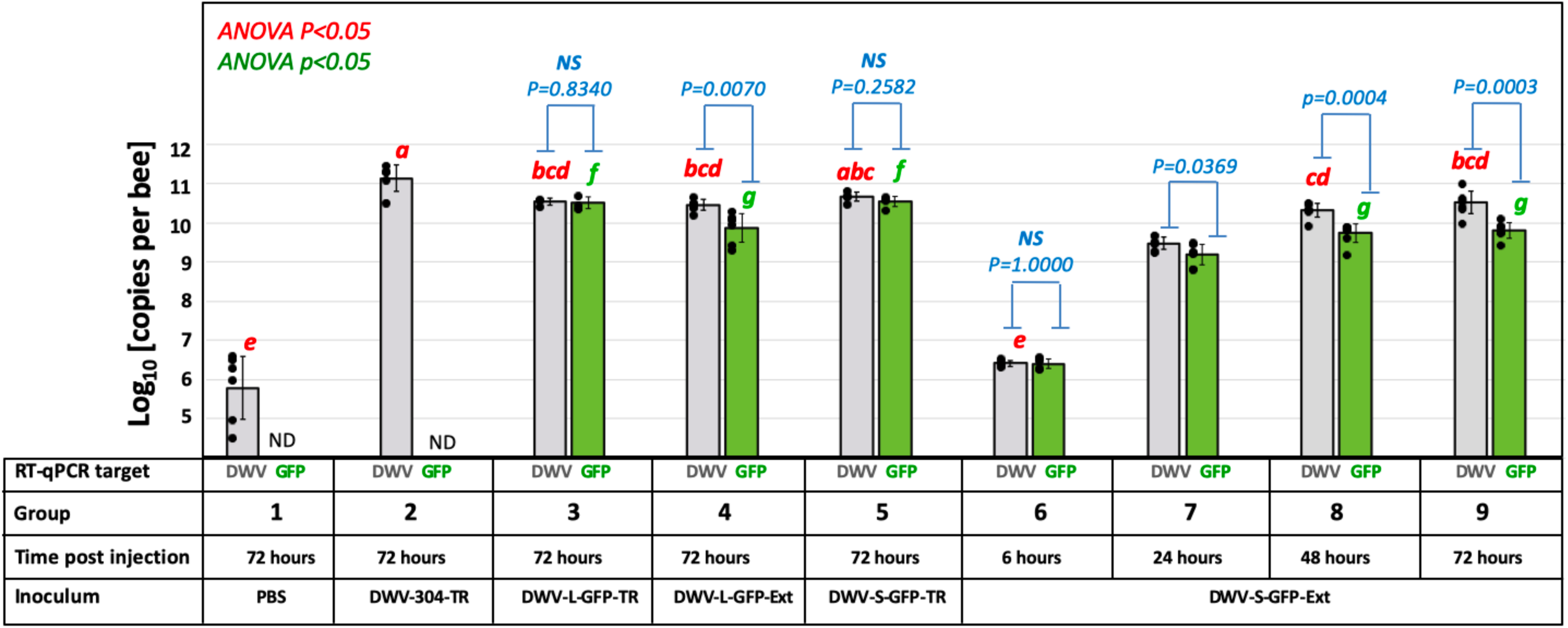
Replication of the GFP-tagged DWV in honey bee pupae. Average copy numbers of DWV and GFP RNA per pupa revealed by RT-qPCR are shown as the light grey and green graphs, respectively, the error bars show ± standard deviation (SD), black dots show DWV RNA or GFP RNA copy numbers in individual pupae. Treatments are shown below the graphs; Inocula: PBS - phosphate buffered saline; suffix -TR, in vitro RNA transcript; suffix -Ext, filtered extract from the pupae infected with the corresponding *in vitro* RNA transcript. Red letters above the bars indicate significantly and non-significantly different groups (ANOVA, P < 0.05). Blue bars – statistical significance of the DWV and GFP copy numbers within the same group NS - non significant, ANOVA P > 0.05).

**Figure 3.**
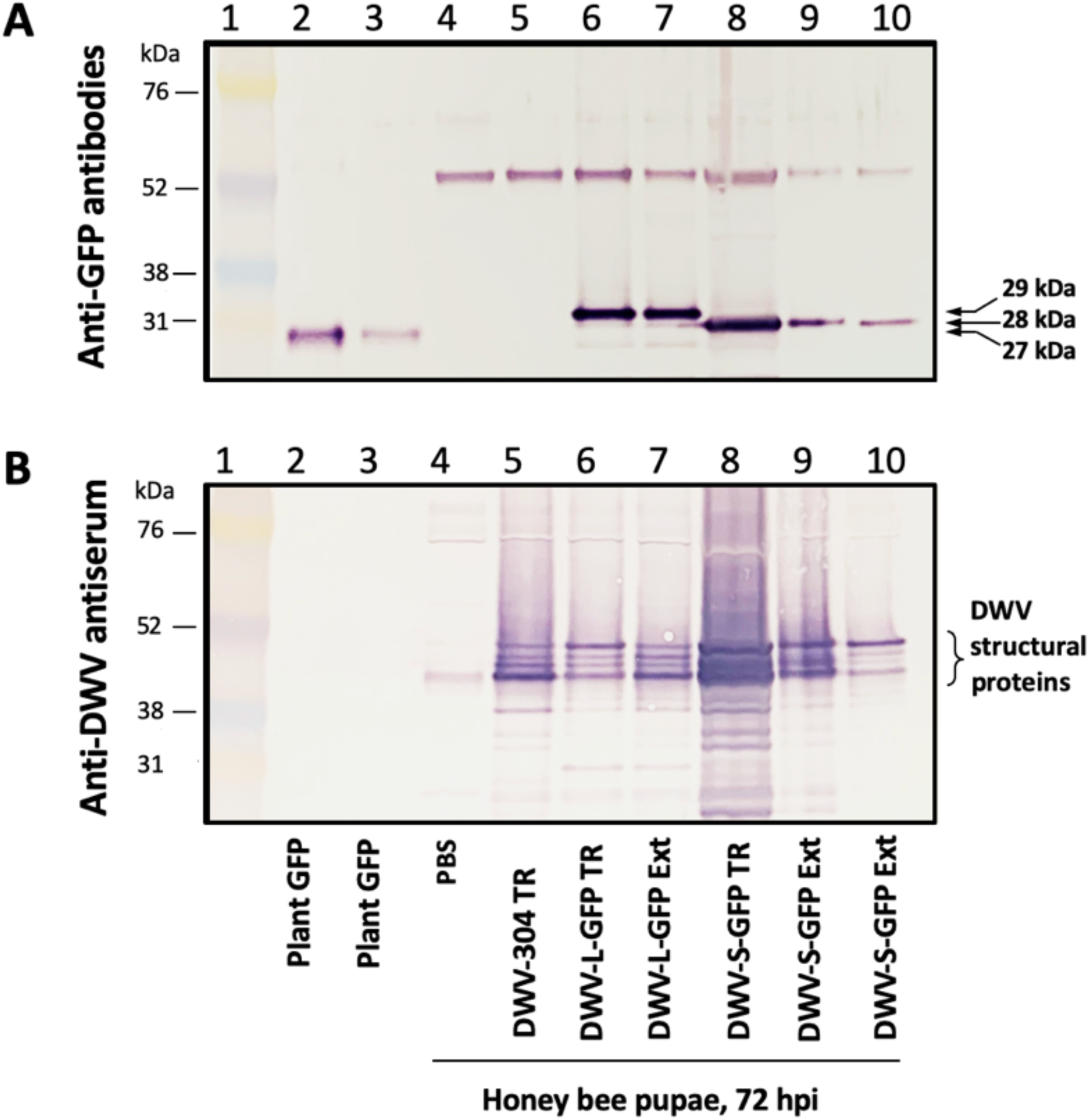
Expression of GFP in honey bee pupae from DWV vector. Western blot analysis using (A) anti-GFP antibodies, and (B) antiserum to DWV virus particles. Protein molecular weight markers (lane 1); lanes 2 and 3, GFP transiently expressed in *Nicotiana benthamiana* plants (lanes 2 and 3); honey bee pupae, 72 hpi, injected with PBS buffer (lane 4), DWV-304 *in vitro* RNA transcript (lane 5); DWV-L-GFP *in vitro* RNA transcript (lane 6); DWV-L-GFP filtered extract (lane 7); DWV-S-GFP in vitro RNA transcript (lane 8); DWV-S-GFP filtered extract (lanes 9 and 10). Expected molecular weights of GFP peptides are shown.

**Figure 4.**
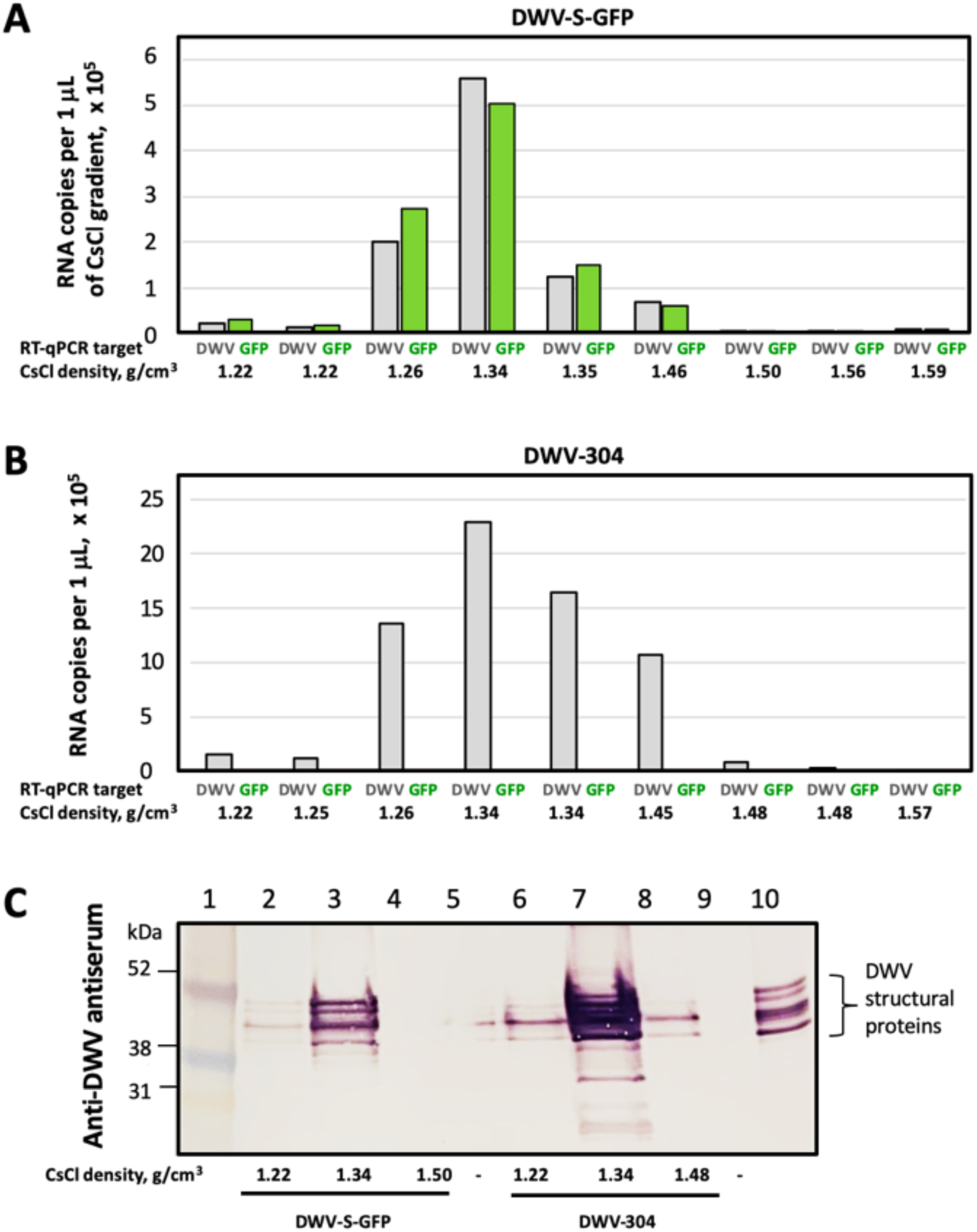
Encapsidation of the DWV-S-GFP genomic RNA. Virus particles accumulated in the honey bee pupae infected with (A) DWV-S-GFP and (B) DWV-304 in vitro RNA transcripts, 72 hpi were separated by CsCl buoyant density gradient centrifugations. The DWV and GFP RNA levels were quantified by RT-qPCR in the density gradient fractions indicated shown below. (C) Western blot analysis of the gradient fractions using antiserum to DWV virus particles. Lane 1, molecular weight standards; lanes 2-4, DWV-S-GFP CsCl gradient fractions; lanes 6-8, DWV-304 CsCl gradient fractions; lane 10, pupae infected with DWV-S-GFP; lanes 5 and 9, empty.

**Figure 5.**
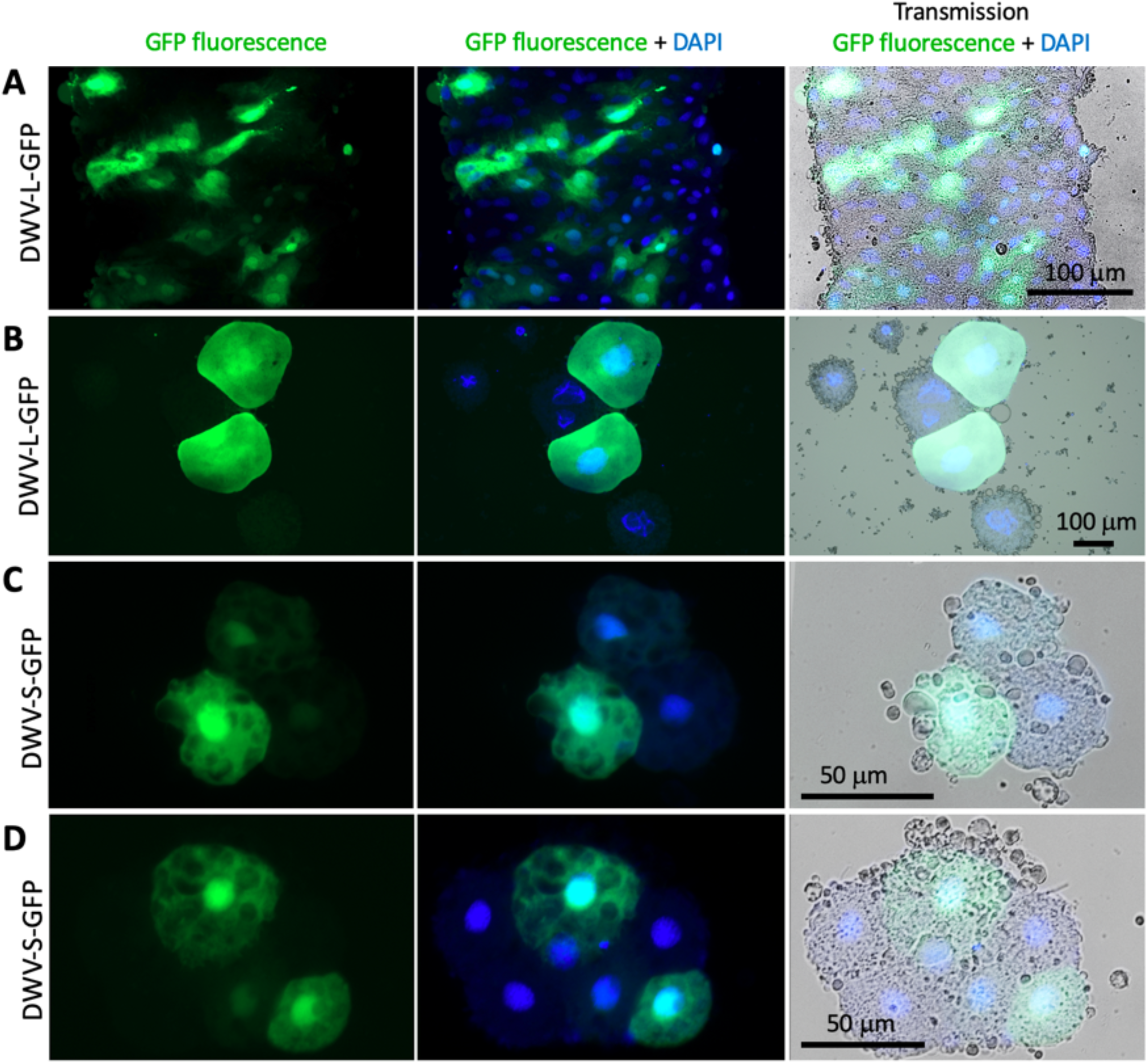
Localization of GFP and DWV structural proteins in the honey bee cells infected with GFP-tagged DWV. Fluorescent microscopy of the cells of the pupae 72 hours post injection with the filtered extracts containing 10^7^ genome equivalents (GE) of DWV-S-GFP (A, B) or DWV-S-GFP (C, D). Left panels, GFP fluorescence (green); middle panels, GFP fluorescence (green) and DAPI nuclear staining (blue); right panels, microscopy transmission images combined with GFP fluorescence and DAPI staining.

**Figure 6.**
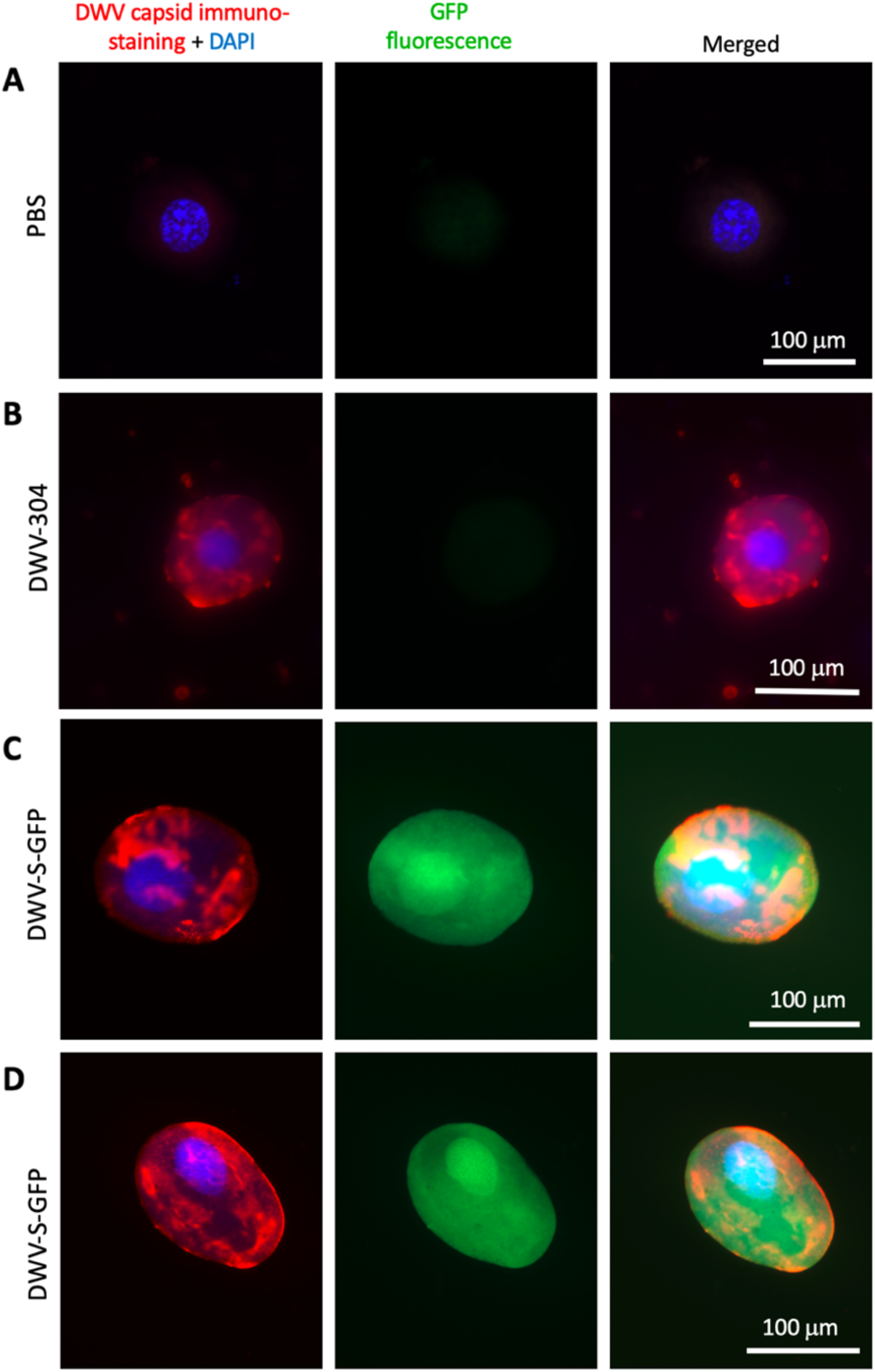
Localization of GFP and DWV structural proteins in the honey bee cells infected with GFP-tagged DWV. Fluorescent microscopy of the cells of the pupae 72 hours post injection with the filtered extracts containing 10^7^ genome equivalents of DWV-304 (B), or DWV-S-GFP (C, D). Left panels, immunostaining of the DWV capsids with anti-DWV antisera, Alexa Fluor 647 (red) and DAPI nuclear staining (blue); middle panels, GFP fluorescence (green); right panel, GFP fluorescence combined with Alexa Fluor 647 and DAPI staining.

**Figure 7.**
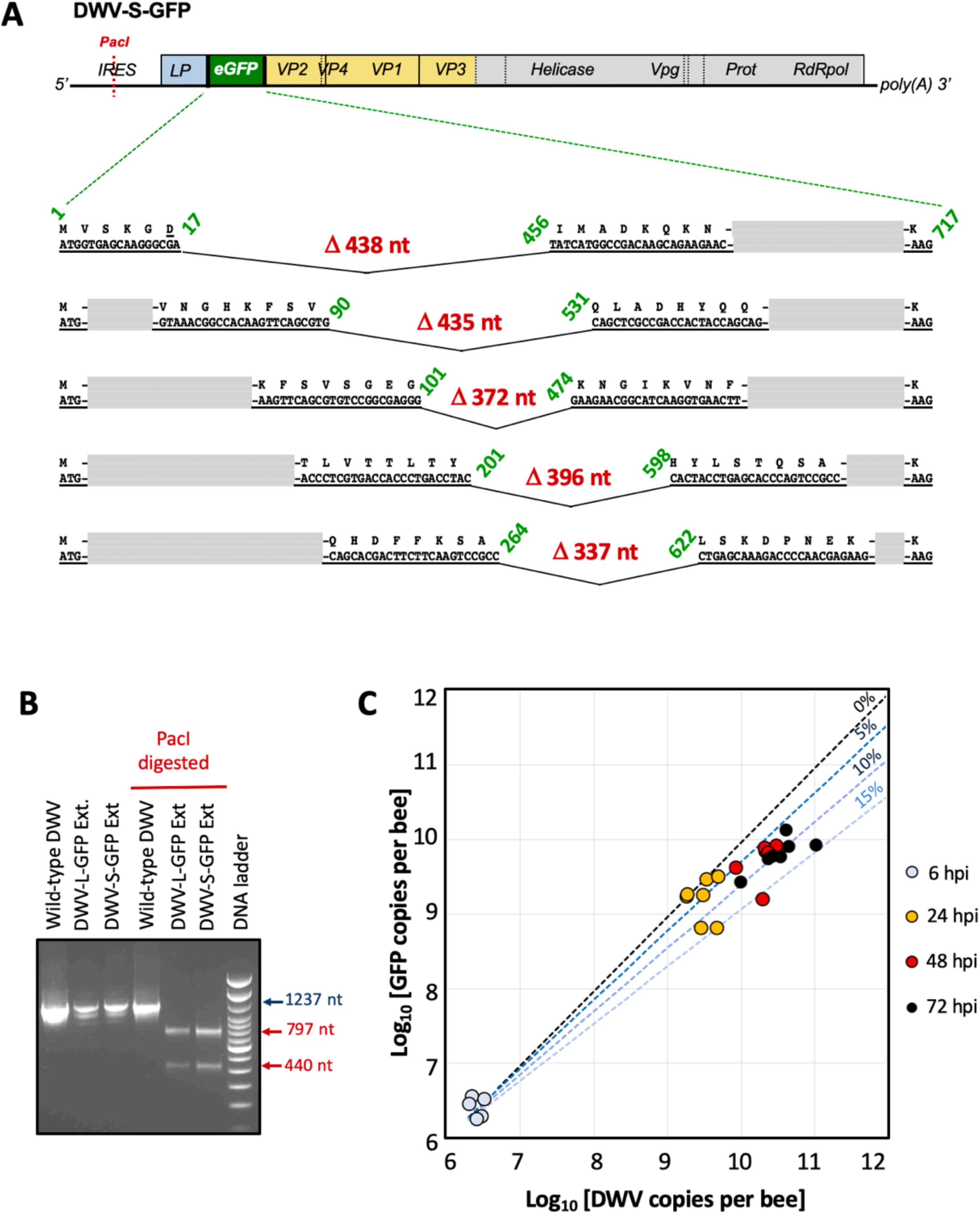
Genetic stability of the GFP-tagged DWV. (A) Deletions of the GFP-coding sequence in the DWV-S-GFP progeny accumulated in a honey bee pupae 72 hours post injection (hpi) with the virus extract. The RT-PCR fragments were generated using the primers flanking GFP insert and cloned into a plasmid vector. Shown are the nucleotide sequences and the corresponding amino acid sequences adjacent to the deletion breakpoints, the numbers indicate the positions of the GFP sequence joined in the deletants and the sizes of deletions. The integrity of the viral ORF was restored in the case of all deletions. (B) Analysis of the 5’ RT-PCR region. The RT-PCR fragments corresponding to the 5’ of DWV were produced using combined samples of the pupae injected with DWV-L-GFP and DWV-S-GFP extracts at 72 hpi, pupae with the wild-type DWV was used as a control. The RT-PCR fragments were digested with *Pac*I restriction enzyme, the site was present only in the cDNA clone-derived DWV. (C) Copy numbers of DWV (X-axis) and GFP (Y-axis) RNA targets revealed by RT-qPCR in individual honey bee pupae 6, 24, 48 and 72 hpi with the virus extracts containing 107 genome equivalents of DWV-S-GFP. Dashed lines indicate projected ratios of DWV and GFP with 0%, 5%, 10% and 15% probability of generation deletion mutant during genome replication cycle starting from the average copy numbers observed at 6 hpi.

### 2.3. RT-qPCR and virus progeny analysis

Total RNA extracted from individual honey bee pupae using Trizol reagent (Ambion) or CsCl density gradient fractions were further purified using a RNeasy kit (QIAGEN) according to manufactures’ instructions. Quantification of the DWV and GFP RNA and actin mRNA loads in individual honey bee pupae and associated *Varroa* mites were quantified by reverse transcription quantitative PCR (RT-qPCR), which included cDNA synthesis using random hexanucleotides. RT-qPCR and cloning of the RT-PCR fragments were carried out essentially as previously described [14], using primers listed in Table S1. DWV RNA copy numbers were log-transformed prior to statistical analyses.

### 2.4. Detection of GFP and DWV structural proteins in honey bees by Western blot and immunofluorescence microscopy

Western blot analysis was carried out using a WesternBreeze kit (Invitrogen) following manufacturer’s instructions. The aliquots of homogenized honey bee pupae protein (50 μg) or 20 μL aliquots of CsCl gradient were heated at 95°C for 10 min in a sodium dodecyl sulphate (SDS) sample buffer (WesternBreeze; Invitrogen) and separated on SDS 4 - 12 % Bis-Tri -polyacrylamide gel electrophoresis gels (Invitrogen). Polyclonal rabbit anti-GFP antibodies (ThermoFisher, Catalogue number PA1-980A) or rabbit polyclonal antisera raised to DWV virus particles purified using CsCl gradient centrifugation [15] were used to detect eGFP and DWV structural proteins of DWV, respectively. The leaf tissue of *Nicotiana benthamiana* agroinfiltrated with the construct expressing 27 kDa monomeric GFP [16], was used as a positive control for GFP. For visualization of eGFP expression in the injected pupae, the pupae at 72 hpi were dissected and hemolymph sample (approximately 30 μL per pupae) with cells and fragments of tissue were collected, fixed with 500 μL of 4% paraformaldehyde in PBS for 5 minutes at room temperature, centrifuged at 2500 rpm in Eppendorf centrifuge for 2 minutes, and washed 3 times with 500 μL of PBS supplemented with 0.05 % Tween-20. For DWV immunostaining, the cells were incubated with rabbit antisera raised against DWV particles {15] overnight at +4°C, the Alexa Fluor 647-labelled goat anti-rabbit antibodies (Invitrogen, Catalogue number A11008) were used. Fluorescent microscopy was carried out using a Zeiss AX10 compound microscope (Zeiss Model Imager M2) fitted with and Axoio.M2 with camera 105. Visualization was carried out as using the following excitation / emission wavelengths: eGFP - 488 nm / 520 nm, DAPI - 359 nm / 461 nm, Alexa Fluor 647 – 651 nm / 676 nm.

## 3. Results

### 3.1. Design of GFP-tagged DWV cDNA constructs

A full-length infectious cDNA clone of a US strain of wild-type DWV type A, pDWV-304 [11], was used to design two recombinant DWV constructs expressing enhanced GFP, pDWV-L-GFP and pDWV-S-GFP (Figure 1, Text S1, Text S2). In the both constructs, the eGFP-coding sequence was inserted between the sequences coding for the leader protein (LP) and the structural virus protein 2 (VP2)-coding sequences in frame to allow translation of ORF with eGFP inserts (Figure 1). The C-terminus of eGFP was linked to the amino acid position “minus 6” of the proposed proteolytic cleavage peptide between the LP and VP2 (VQAKPEMDNPNPG) which is tentatively targeted by DWV 3C protease (10). The N-terminus of eGFP was fused to the LP in two configurations: either via a proposed full proteolytic cleavage peptide VQAKPEMDNPNPG present at the LP-VP2 border in the clone pDWV-L-GFP, where proteolysis takes place between the glutamate and methionine residues [10;17], or via a shorter peptide, VQAKPE, corresponding to the proposed C-terminus of the LP after proteolysis [10,17] (Figure 1). In the case of the pDWV-S-GFP construct, the linking peptide did not include the complete proposed proteolytic site and therefore this construct was expected to express LP-GFP as a fusion protein. In both *egfp-*tagged DWV constructs, proteolytic processing of the viral polypeptide between eGFP and VP2 was expected to take place, ensuring the generation of wild-type structural viral proteins required for encapsidation of DWV RNA.

### 3.2. Infectivity the cDNA clone-derived GFP-tagged DWV in honey bee pupae

Cell cultures are commonly used for studying the clone-derived RNA viruses of vertebrates and invertebrates, but this option is not readily applied to cloned DWV. Honey bee cell cultures suitable for work with DWV are not readily available. A recently reported honeybee cell line is infected already with DWV [18] and no other existing invertebrate cell cultures support DWV replication. Therefore, we injected honey bee pupae with viral *in vitro* RNA transcripts in order to recover and study recombinant clone-derived DWV, as reported previously [11]. *In vitro* RNA transcripts from the constructs pDWV-L-GFP and pDWV-S-GFP were infectious when injected into honey bee pupae intra-abdominally. Quantification of *egfp* target by RT-qPCR (Fig. 2, Groups 3 and 5, GFP; Table S2) showed that at 72 hours post injection (hpi) the pupae injected with pDWV-L-GFP and pDWV-S-GFP had high statistically identical (P = 0.802, df = 6, ANOVA; Table S3) levels of genomic RNAs of DWV-L-GFP, ranging from 10.37 to 10.72 log_10_ genome equivalents (GE) per pupa, and DWV-S-GFP, ranging from 10.33 to 10.66 log_10_ GE per pupa (Figure 1, Groups 3 and 5; Table S2). Levels of the in vitro transcript-derived *egfp*-tagged viruses at 72 dpi were far higher than those previously shown for of mutagenized non-replicating DWV in vitro transcripts [11] and control buffer injection (Figure 2, Group 1) ca. 5 to 7 log_10_. Importantly, RT-qPCR revealed that the copy numbers of the *egfp* target (quantification of the genomes with the *egfp* insert only) and the DWV target (quantification of all DWV genomes, with and without *egfp)* were not significantly different in the pupae injected with the recombinant *egfp*-tagged DWV transcripts (Figure. 2; Group 3: P = 0.834, df = 5; Group 5: P = 0.258, df = 7; ANOVA; Table S3). This showed that most of the recovered virus genomes in the pupae injected with the DWV-L-GFP and DWV-S-GFP *in vitro* transcripts retained *egfp*, and the presence of 717 nt *egfp* insert (approximately 7 % of the DWV RNA) between the genes for LP and VP2 did not prevent replication of the modified viruses, *i.e.*, by disrupting potential RNA elements essential for virus RNA replication [19].

Quantification of the DWV target showed that in the pupae injected with non-modified DWV *in vitro* RNA transcript from pDWV-304 the virus replicated to high levels (range: 10.51 to 11.47 log_10_ GE/pupa; 11.14 ± 0.338 log_10_ GE/pupa, mean ± SD, by 72 hpi; Table S2), which were significantly higher (4-fold) than in the DWV-L-GFP transcript-injected insects (P = 0.0398, df = 7, ANOVA; Figure 2, Groups 2 and 3). Compared to the DWV-S-GFP transcript injection group, the levels of unmodified DWV-304 were 3-fold higher, though this difference was not significant (P = 0.0505, df = 8, ANOVA; Figure 2, Groups 2 and 5). The levels of DWV in the control phosphate buffered saline (PBS)-injected pupae (Figure 2, Group 1, range: 4.48 to 6.59 log_10_ GE/pupa; 5.79 ± 0.801 log_10_ GE/pupa; mean ± SD) were significantly lower (tens of thousands times) compared to those in any of the transcript-injected groups (Figure 2; Table S3, P<0.0001, ANOVA). As expected, the *egfp* RNA was not detected in the buffer control (PBS)- and the DWV-304-injected bees (Figure 2, Groups 1 and 2). Reduced levels of *egfp*-tagged DWV compared to the unmodified DWV-304, could reflect a fitness cost to the virus from increasing the genome size by 7 % with a non-essential sequence.

To test whether expression of eGFP occurred in the pupae injected with the *in vitro* transcripts from the *egfp*-tagged DWV constructs, at 3 dpi, the pupal extracts were subjected to Western blot analysis with anti-GFP antibodies (Figure 3A). Control pupae injected with the buffer (PBS) or the wild-type DWV transcript (Figure 3A, lanes 4, 5), also at 72 hpi, were also included alongside the extracts from plant (*Nicotiana benthamiana*) tissues expressing monomeric GFP of 27 kDa (Figure 3A, Lanes 2, 3) [16].

The Western blot analysis with anti-GFP antibodies clearly showed accumulation of the 29 kDa GFP protein in pupae injected with the DWV-L-GFP transcript, indicating accumulation of fully processed free eGFP. Surprisingly, in the case of DWV-S-GFP transcript accumulation of the 28 kDa GFP rather than LP-GFP fusion was observed (Figure 3A, lane 8), indicating that proteolytic cleavage of the shorter version of the LP-GFP linking peptide also took place. It is possible that the LP-GFP linking peptide in the case of DWV-S-GFP led to cleavage between glutamate and glycine residues of the peptide VQAKPEGAP, the glutamate-glycine cleavage site was demonstrated for the DWV peptide linking VP1 and VP3 [10,17]. The observed differences of molecular weight of DWV vectors compared to free monomeric GFP in the plant tissue controls (28 kDa for DWV-S-GFP, 29 kDa for DWV-L-GFP, and 27 kDa for plant-expressed free GFP) was the result of the residual peptides left after the proteolytic cleavage of the eGFP from the viral polyprotein and was in a good agreement with their molecular weight predictions of 27.97 kDa, 28.69 kDa, and 26.93 kDa, respectively (Figure S1). Importantly, the Western blot analysis with anti-GFP antibodies showed the absence of detectable quantities of polypeptides heavier than the expected monomeric eGFP, which could be a result of the accumulation of the eGFP fusions with LP and structural VP2 proteins (Figure 3A). The visible band with molecular weight about 55 kDa was present not only in the case of DWV-S-GFP and DWV-S-GFP, but also in the case of the PBS-injected pupae with low DWV levels DWV-free and the DWV-304 injected with high DWV levels (Figure 3A,B). This indicated complete proteolytic processing of the GFP-VP2 peptide in the case of both DWV-L-GFP and DWV-S-GFP (Figure 3A) suggesting that maturation of the structural proteins capable of packaging DWV genomic RNA took place during replication of the *egfp*-tagged DWV.

### 3.3. Analysis of encapsidation of the egfp-tagged DWV RNA genomes

DWV genomic RNAs with the *egfp* insert were approximately 7 % longer than wild-type DWV RNA (Text S2) potentially increasing the length of the modified genomic RNA beyond the RNA size limit for encapsidation. In addition, the insertion of a foreign sequence might potentially disrupt yet unknown RNA signals required for the formation of stable virus particles [19,20]. Therefore, it was necessary to determine if stable virus particles were formed in honey bee pupae infected with the *egfp*-tagged DWV and to compare them with wild-type DWV particles. To do this, we carried out cesium chloride (CsCl) gradient centrifugation of the filtered extracts from honey bee pupae 72 hpi with the transcripts from the cDNA clones DWV-S-GFP and DWV-304 (unmodified DWV). Following ultracentrifugation, the fractions with densities ranging from 1.22 g/cm^3^ to 1.58 g/cm^3^ were collected and the copy numbers of DWV and GFP RNA targets were quantified for each fraction by RT-qPCR (Figure 4A,B).

We found that in the case of DWV-S-GFP, the 1.34 g/cm^3^ buoyant density fraction showed the highest levels of both GFP and DWV targets (Figure 4A). In the case of DWV-304, the highest level of DWV RNA genome was also in the same 1.34 g/cm^3^ CsCl gradient fraction (Figure 4B), which was within the range of the predicted buoyant density for viruses of the family Iflaviridae in CsCl (1.29 - 1.38 g/cm3) [21]. In addition, Western blot analysis with the antisera against DWV virus particles confirmed the highest load of the DWV capsid proteins in the fractions with the highest levels of DWV for both DWV-304 and DWV-S-GFP (Figure 4C). These results showed that the DWV RNA genome with a 750 nt insertion (717 nt *egfp* gene and the sequences coding for duplicated flanking peptides) was encapsidated into particles with the stability and buoyant density similar to those of unmodified DWV.

Encapsidation of DWV genomes with *egfp* inserts, both DWV-L-GFP and DWV-S-GFP, was further confirmed by infectivity of the filtered tissue extracts prepared from the transcript-infected honeybee pupae sourced at 72 hpi. Extracts selected for injection had nearly equal copy numbers of DWV and GFP RNA by RT-qPCR (Figure 2, Groups 3 and 5), indicating that they mostly contained the recombinant DWV genomes with the *egfp* inserts. In the case of DWV-S-GFP, an extract from the same transcript-infected pupae was used in CsCl buoyant density analysis (Figure 4A). Honey bee pupae injected with doses of 7 log_10_ GE of DWV-L-GFP or DWV-S-GFP showed development of recombinant virus infections. According to GFP quantification at 72 hpi, the levels of DWV-L-GFP ranged from 9.31 to 10.29 log_10_ GE/pupa (9.87 ± 0.364 log10 GE/pupa; mean ± SD), and the levels of DWV-S-GFP ranged from 9.42 to 10.12 log_10_ GE/pupa (9.81 ± 0.200 log10 GE/pupa; mean ± SD), Figure 2, Groups 5 and 10. There were no significant differences between the levels of DWV-L-GFP and DWV-S-GFP (P = 0.275, dF = 12; Table S3).

We further confirmed replication of recombinant DWV genotypes with intact *egfp* inserts in the pupae injected with DWV-L-GFP and DWV-S-GFP-derived extracts by Western blot analysis with the anti-GFP antibodies. Specifically, at 72 hpi monomeric eGFP bands of the same size were observed in both the extract-injected pupae and in the pupae injected with corresponding transcripts (Figure 3A, DWV-L-GFP: lanes 7, 8; DWV-S-GFP; lanes 8-10).

### 3.4. Expression of eGFP reporter from the DWV RNA vectors

Fluorescence microscopy was used to assess eGFP expression in honey bee pupae injected with the DWV-L-GFP and DWV-S-GFP filtered extracts at 72 dpi. Individual hemolymph cells and tissue fragments, possibly fragments of immature intestinal epithelium, of the pupae injected with DWV-L-GFP (Figure 5A,B) and DWV-S-GFP (Figure 5C,D; Figure 6C,D) showed strong GFP fluorescence, which was not observed in the case of buffer (PBS) and the unmodified DWV (DWV-304) injections (Figure 6A,B).

The GFP fluorescence in the case of the *egfp*-tagged DWV variants was observed in both cytoplasm and nucleoplasm as confirmed by co-localization of the GFP and DAPI fluorescence (Figure 6C,D), which was in agreement with previously reported localization of free GFP in insect cells [22]. Such localization of eGFP expressed from DWV vectors confirmed our Western blot analysis findings (Figure 3A) that complete proteolytic excision of eGFP from the viral polyprotein occurred. Fluorescent microscopy of the groups of cells in the case of DWV-L-GFP and DWV-S-GFP injections showed presence of these cells with different levels of GFP expression in the same tissue fragment, which was evident from the DAPI fluorescence and transmission light microscopy (Figure 5). We also demonstrate, by using immunofluorescent microscopy with antibodies against the DWV capsid, that GFP was expressed in DWV-infected cells (Figure 6C,D). Immunofluorescence microscopy revealed that DWV capsid proteins were forming aggregates in the cytoplasm, while GFP was diffusely distributed in both cytoplasm and nucleoplasm in the same cells (Figure 6C,D).

### 3.5. Genetic stability of egfp-tagged DWV RNA

Quantification of the eGFP and DWV RNA targets in pupae injected with extracts containing 10^7^ GE of recovered DWV-L-GFP or DWV-S-GFP showed that at 72 dpi DWV copy numbers were significantly higher (P < 0.05, ANOVA) than those of GFP indicating accumulation of DWV variants without *egfp* insert. Both *egfp*-tagged DWV variants showed similar levels of GFP and DWV at 72 hpi (Figure 2, Groups 4 and 9, GFP: P = 0. 0.715393, dF = 12; DWV: P = 0.663399, dF = 11, ANOVA), and similar difference between DWV and GFP copy numbers in individual pupae (Figure S2, Groups 4 and 9, P = 0.1578, dF = 12, ANOVA). In the case of DWV-S-GFP injection, pupae were also sampled at 6, 24, and 48 hpi, showing non-significant differences between DWV and GFP copy numbers within each time group at 6 and 24 hpi, while at 48 dpi the copy number of DWV RNA significantly exceeded that of the GFP RNA (Figure 2, Groups 6, 7, 8). Such differences suggest that in addition to the *egfp*-tagged DWV, injected pupae contained DWV without these inserts.

To determine the nature of the deletions in the DWV-S-GFP progeny, we amplified the section which included to the *egfp* insert by RT-PCR using the primers flanking *egfp* insert (Table S1) in the pooled DWV-S-GFP extract-injected pupae at 72 hpi and cloned the heterologous fragments with *egfp* deletions into a plasmid vector. A sample of clones with inserts of different sizes was sequenced and we found different deletions of the 717 nt eGFP-coding sequence ranging from 357 to 438 nt (Figure 7A), Notably, in the case of all deletions, the ORF was restored which was essential for translation of DWV RNA.

We tested whether the accumulation of DWV variants without the *egfp* insert was a result of amplification of background DWV present in the recipient pupae at low levels (Figure 2, Group 1) or the accumulation of the DWV-L-GFP and DWV-S-GFP progeny with deletion of in the *egfp* gene. To do this, we determined if the *Pac*I restriction site (Figure 1) was present in the 5’ IRES of the DWV progeny. The *Pac*I sequence was absent in the wild-type DWV strains, but was introduced to the 5’ IRES sequence of the parental DWV-304 clone [11] and was retained in both DWV-L-GFP and DWV-S-GFP, therefore it could be used as a genetic marker to distinguish between wild-type DWV and cDNA clone-derived DWV. We demonstrated complete *Pac*I digestion of the RT-PCR fragments amplified using samples of 72 hpi pupa injected with DWV-S-GFP and DWV-L-GFP inocula, but not with the wild-type DWV (Figure 7B), which indicated that all DWV genomes, including those without the *egfp* insert, were clone-derived.

When DWV and GFP GE number were plotted on a 2-dimensional graph (Figure 7C, X axis – log_10_ DWV GE per pupae; Y axis – log_10_ GFP GE per pupae), it became obvious that the proportion of the mutants with deletions of *egfp* was increasing with time. We estimate that the observed rate of accumulation in progeny with *egfp* deletions could be explained by an approximately 10 % chance of such a deletion event per RNA replication event (Fig. 7 C, dotted projection lines). Notably, only deletion events within *egfp* inserts which resulted in generation of restored viral ORF were viable were fixed in the viral population. Despite accumulation of such deletion mutants, RT-qPCR results suggest that between 6 hpi and 72 hpi the load of the *egfp*-tagged virus had increased on average approximately 2500 times, ranging from 1030 to 5196 times in individual pupa, compared to average GE counts at 6 hpi (Figure 2, Groups 6 and 9; Fig. 7C, 6 hpi *vs* 72 hpi), clearly indicating strong infectivity of DWV genomes with the inserted foreign sequence (Figure 1). These results are in a good agreement with detection of GFP expression by fluorescent microscopy of the recombinant DWV extract-infected pupae (Figures 5 and 6).

## 4. Discussion

In this study we designed and evaluated DWV-based virus vectors, the first constructed using the genome of the RNA virus naturally infecting honey bees [10], and demonstrated their ability to achieve high levels of expression of the eGFP reporter in honey bees. DWV is the major virus pathogen of honey bees and is associated with increased colony losses worldwide that have affected pollination services and food security [23,24]. DWV belongs to the family Iflaviridae (order Picornavirales) which includes 15 recognized and 18 pending virus species infecting a wide range of insects [21]. It is very likely that the real diversity and distribution of this virus group is much wider and therefore it is likely to have a great impact on many invertebrate species. For example, among 1,445 novel viruses discovered in a metagenomic analysis of viromes of 220 invertebrate species [25] approximately 400 of them belong to Picorna-Calici clade, some of which are likely to be the members of the order Picornavirales, including approximately 50 showing highest homology with the previously identified iflaviruses. Despite a growing number of sequenced RNA viruses in invertebrates, there are significant gaps in the knowledge of their biology and interactions with host(s). Therefore, the use of novel DWV vector constructs, in particular those expressing fluorescent reporters (e. g. eGFP) allowing non-invasive monitoring of the infection development in live cells and tissues, could provide novel insights into many viruses infecting invertebrates.

We demonstrated that DWV RNA genome could be used to express a 29 kD foreign protein (eGFP) when the *egfp* gene was inserted into the viral ORF in frame between the sequences coding for the LP and VP2 structural protein. The eGFP polypeptide was flanked with the peptides derived from the DWV LP-VP2 interface allowing release of free eGFP following the cleavage by DWV 3C protease (Figure 1). The expression eGFP was demonstrated by observation of GFP fluorescence in the DWV-L-GFP- and DWV-S-GFP-infected cells of honey bees by fluorescent microscopy (Figures 5 and 6), and by the detection of polypeptides of expected sizes in Western blots experiments using antibodies against GFP (Figure 3A). The recombinant DWV genomes replicated to high levels, approximately 10^10^ to 10^11^ GE per pupae by 72 hpi (Figure 2), which indicated that insertion of the foreign gene did not disrupt any cis-RNA elements required for replication [19].

There was a possibility, that increase of DWV genomic RNA by 7 % as a result of *egfp* insertion could affect encapsidation of the viral RNA. It was previously demonstrated that an RNA size limit for encapsidation exists for single stranded RNA viruses with icosahedral symmetry of virus particles structurally resembling those of Iflaviruses, such as Turnip crinkle virus [26,27]. Also, potential RNA elements required for DWV RNA encapsidation could be disrupted [19,20].

Therefore, we used CsCl density gradient centrifugation to test if formation of stable DWV particles containing DWV genomic RNA with *egfp* inserts took place in in the pupae injected with the DWV-S-GFP and DWV-L-GFP in vitro RNA transcripts. We found that DWV genome with *egfp* insertion was packaged into stable virus particles and had the same physical properties as those containing unmodified DWV RNA (Figure 4).

One of the aims of this study was to develop a tagged DWV vector, which could be used for tracking virus replication in the cells and tissues of the DWV host(s). Indeed, we observed bright GFP fluorescence in the cells of the pupae infected with the extracts containing recovered DWV-L-GFP or DWV-S-GFP. Different cell-types showed GFP fluorescence, which was consistent with previously reported presence of DWV throughout honey bee body in various cell types [28]. Importantly, GFP fluorescence was observed in the pupae infected with the filtered tissue extracts, showing suitability of using DWV vector for large scale studies. In our experiment we found that the amount of the recovered *egfp*-expressing recombinant virus inoculum generated in a single *in vitro* RNA transcript-injected pupae was sufficient for hundreds of pupal injections. Therefore, the eGFP-tagged DWV (USA strain of DWV type A) designed in this study, and other cloned DWV-like viruses [29-31], which could also be tagged with a reporter gene in the way developed in this study. Such tagged DWV-like viruses could be an excellent tool to advance understanding of the mechanisms of vectoring of DWV-like viruses by the *Varroa* mites [32] and the host range of DWV-like viruses in different bee species. Currently, there are conflicting reports on whether DWV-like viruses replicate in their *Varroa* mite vector [33-36] and the use of cloned DWV-like viruses expressing *egfp* during their replication would allow to determine if replication of any variants of DWV-like viruses occur in their *Varroa* mite vector and pinpoint the Varroa mite cells where replication could take place.

We also analyzed of genetic stability of our *egfp* tagged DWV constructs. While deletion of foreign gene from RNA virus vector genomes were reported previously [37,38], this was the first study of the genetic stability of a picorna-like RNA virus (Iflavirus) vector conducted in insects at organism level. Our finding supports previous reports that the size of foreign insert dramatically affects genetic stability of the recombinant virus therefore shorter inserts were more tolerated. For example, for a poliovirus-based vector (approximately 7,500 nt RNA genome), the constructs with inserts with a size below 282 nt were more stable than those with 402 nt or 744 nt [37]. Interestingly, we found that some of the *egfp* deletion mutant genomes generated at 72 hpi in the course of DWV-S-GFP infection in the extract-injected bees still had 200-300 nt-long parts of the eGFP-coding sequence (Figure 7A). It is likely that deletion of non-essential gene insertions, even if these were not interfering with virus replication or encapsidation, could be driven by higher competitiveness of a shorter RNA genome. Moreover, infection of an RNA virus in the host transgenically expressing one of the virus proteins rapidly results in generation of the mutants of this virus with deletion of the gene in the virus genome corresponding to the transgene, as it was demonstrated for infection of Tobacco etch virus (TEV) gene in the plants expressing TEV NIb replicase gene [39]. Longer size of genomic RNAs of DWV-S-GFP, DWV-L-GFP and even their deletion mutants (some of which still contained parts of GFP) might explain their reduced levels compared to parental non-modified DWV-304 at 72 hpi (Figure 2, Groups 2, *vs* Groups 3, 4, 5, 9; DWV probes).

It should be emphasized that although accumulation of the progeny with the deletion of non-essential *egfp* took place, this process was gradual and did not happen immediately. We showed that the *egfp*-tagged viruses replicated in the injected pupae increased their loads thousands of times (Figure 2, Figure 7C) suggesting that even a non-essential gene in an RNA virus genome could be relatively stable (Figure 7C). It is a widely accepted view, that gene exchanges between different virus groups and incorporation of host mRNA or their fragments occur during the RNA virus evolution [40,41]. It is necessary for these foreign inserts to be present in genomic RNA undergoing several rounds of replication events to allow further adjustment to the viral RNA genome context via mutagenesis. Our results suggest that even the foreign gene which did not bring benefits to the virus could be a part of an RNA genome for a prolonged time allowing possibility of mutagenesis, provided that such insertion did not dramatically affect replication or other essential virus functions.

## 5. Conclusions

In this study, the DWV vectors were constructed using a full-length infectious cDNA clone of DWV [11] by inserting sequences coding for the *egfp* reporter gene using a novel arrangement which has not been applied in constructing picorna-like vectors previously [2], and included the insertion of a foreign gene into DWV ORF between the sequences coding for the leader protein (LP) and the structural virus protein (VP2). This novel arrangement of the foreign gene insertion did not disrupt replication or encapsidation. While we have shown the value of this approach for the expression of reporter protein, the same approach might be applied for expressing other proteins or RNA molecules important for functional studies or even for maintaining honey bee health. Indeed, the potential of genetically modified bacteria for providing disease-modulating dsRNA for honey bees has been recently shown [42], and it is quite possible that DWV-based vectors might be enlisted for the same role.

## Supplementary Materials

**Text S1.** Design of pDWV-L-GPF and DWV-S-GFP.

**Text S2:** Nucleotide sequences of the Deformed wing virus vector cDNA clones pDWV-L-GFP and pDWV-S-GFP.

## Author Contributions

For research articles with several authors, a short paragraph specifying their individual contributions must be provided. The following statements should be used “Conceptualization, E.V.R.; methodology, E.V.R., M.C.H., F.P.-F., R.L.H., J.D.E.; validation, E.V.R., J.D.E.; formal analysis, E.V.R.; investigation, E.V.R., K.C., M.C..H., F.P.-F.; resources, E.V.R., F.P.-F., R.L.H., J.D.E.; data curation, E.V.R.; writing—original draft preparation, E.V.R., J.D.E.; writing—review and editing, E.V.R., K.C., M.C.H., F.P.-F., R.L.H., Y.C., J.D.E.; visualization, E.V.R., K.C., R.L.H.; supervision, E.V.R, Y.C., J.D.E.; project administration, J.D.E., Y.C., E.V.R.; funding acquisition, E.V.R., Y.C., J.D.E. All authors have read and agreed to the published version of the manuscript.

## Acknowledgments

We thank Dr. Mike Simone-Finstrom (USDA Baton Rouge, Louisiana) for providing honey bees, Dr. Svetlana Folimonova (University of Florida) for GFP control, and Mr. Zachary Lamas for the symptomatic honeybees infected with wild-type DWV.

---

**Figure S1.**
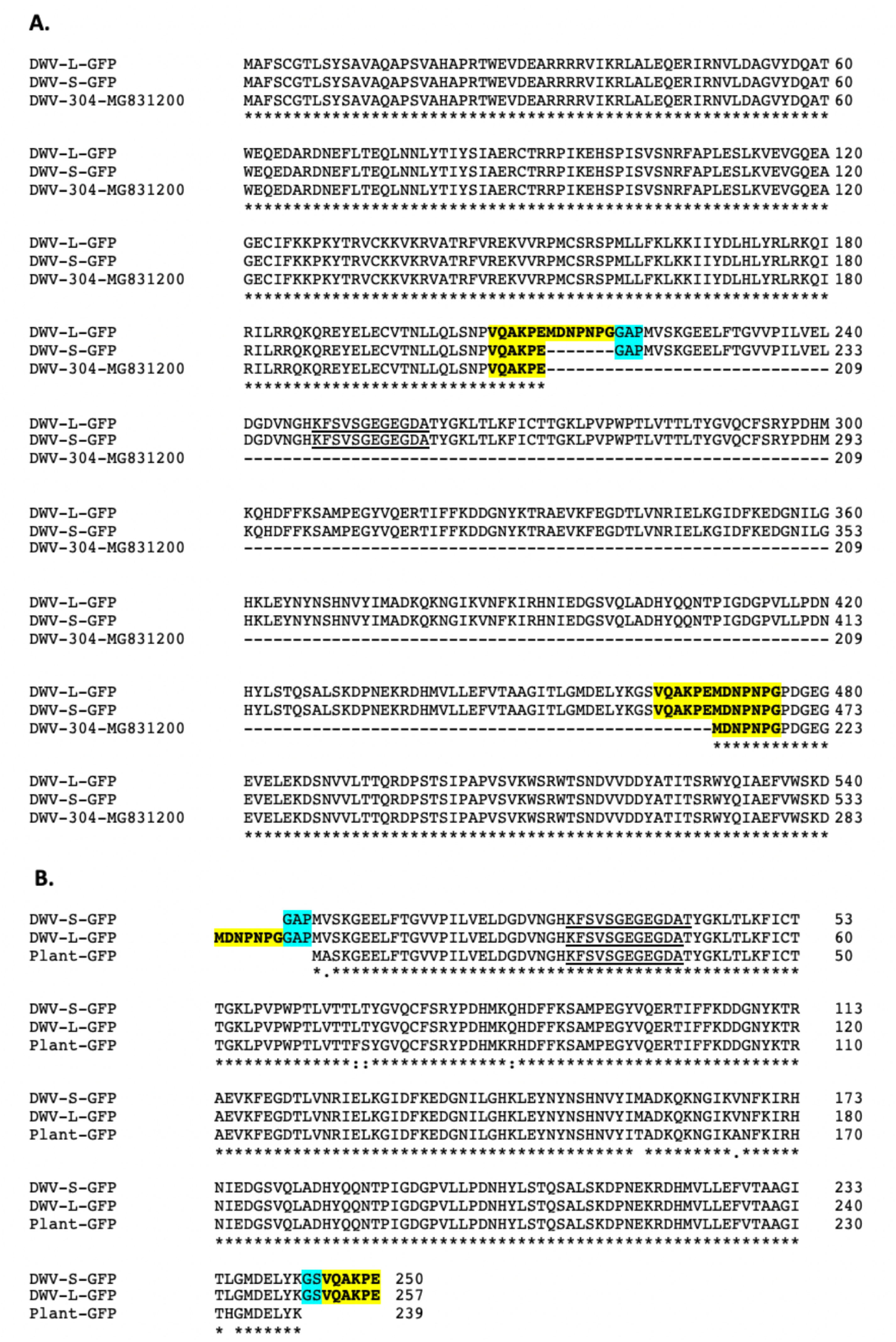
The eGFP-coding sequences inserts in DWV vector. **(A)** Alignment of the N-terminal sections of the polyproteins encoded by infectious cDNA constructs DWV-L-GFP, DWV-S-GFP, and DWV-304 (GenBank accession number MG831200). **(B)** Alignment of the eGFP peptides generated in the honey bee pupae by the DWV 3C proteolytic cleavage of the polyproteins of DWV-S-GFP (27.97 kDa) and DWV-L-GFP (28.69 kDa), and free GFP control expressed in plants (26.93 kDa). Peptide recognized by polyclonal anti-GFP antibodies is underscored. Yellow highlight – parts the DWV LP-VP1 proteolytic cleavage peptide. Blue highlight – amino acid residues introduced by insertion of *Asc*I and *Bam*HI restriction sites.

**Figure S2.**
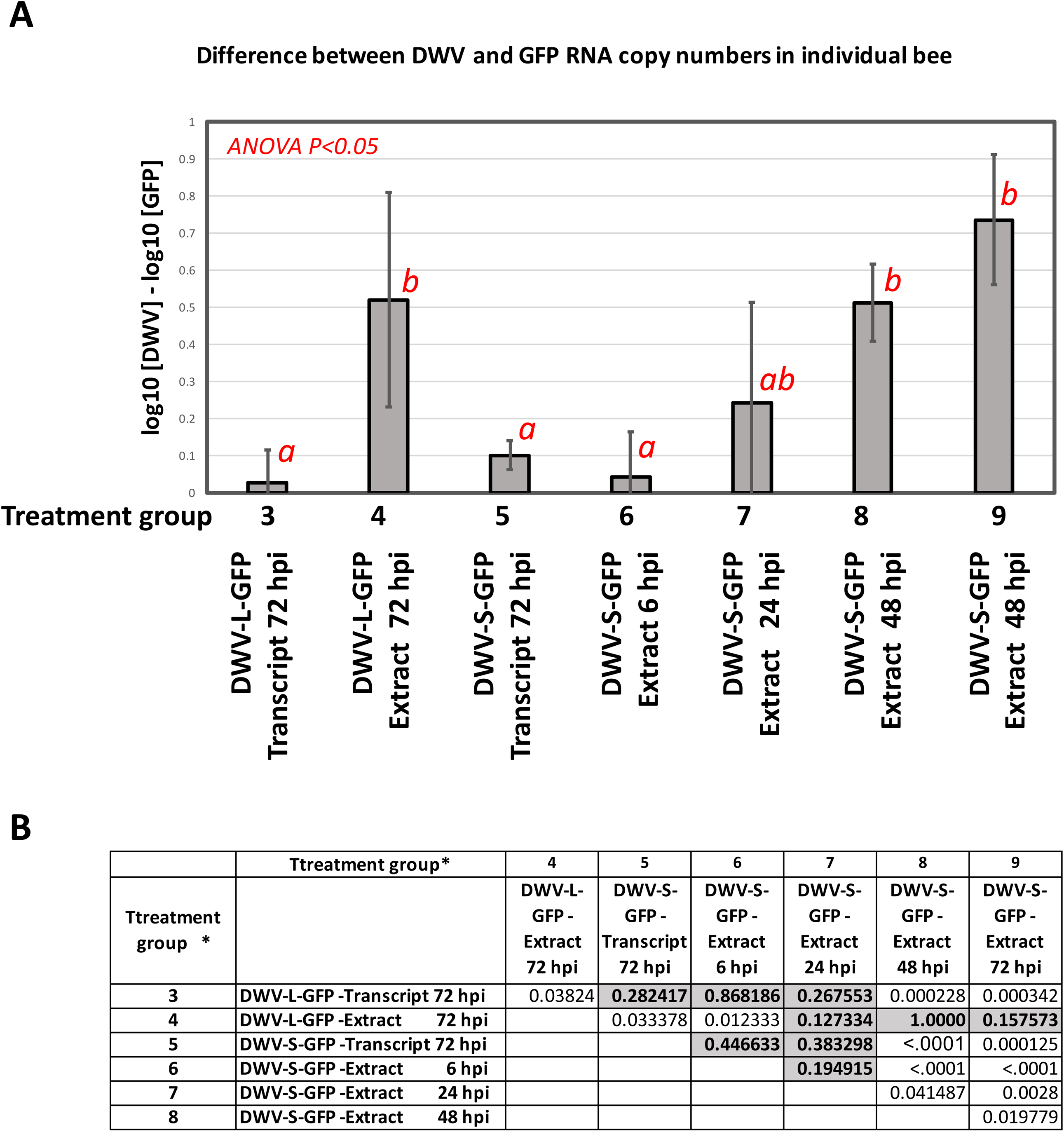
Accumulation of the *egfp* deletant genomes in individual honey bee pupae. (A) Average copy numbers for the differences between DWV and GFP loads (Log_10_(DWV) minus Log_10_(GFP) calculated for individual pupa in the treatment groups are shown as grey graphs, the error bars show ± standard deviation (SD). Treatments are shown below the graphs. Inocula: PBS - phosphate buffered saline; “Transcript”, *in vitro* RNA transcript; Extract, filtered extract from the pupae infected with the corresponding RNA transcript. Red letters above the bars indicate significantly groups (ANOVA, P < 0.05). (B). Statistical significance of the Log_10_(DWV) minus Log_10_(GFP) differences between the treatment groups. P-values, ANOVA analysis. Non-significant co (P>0.05%, ANOVA) are highlighted

**Table S1.**
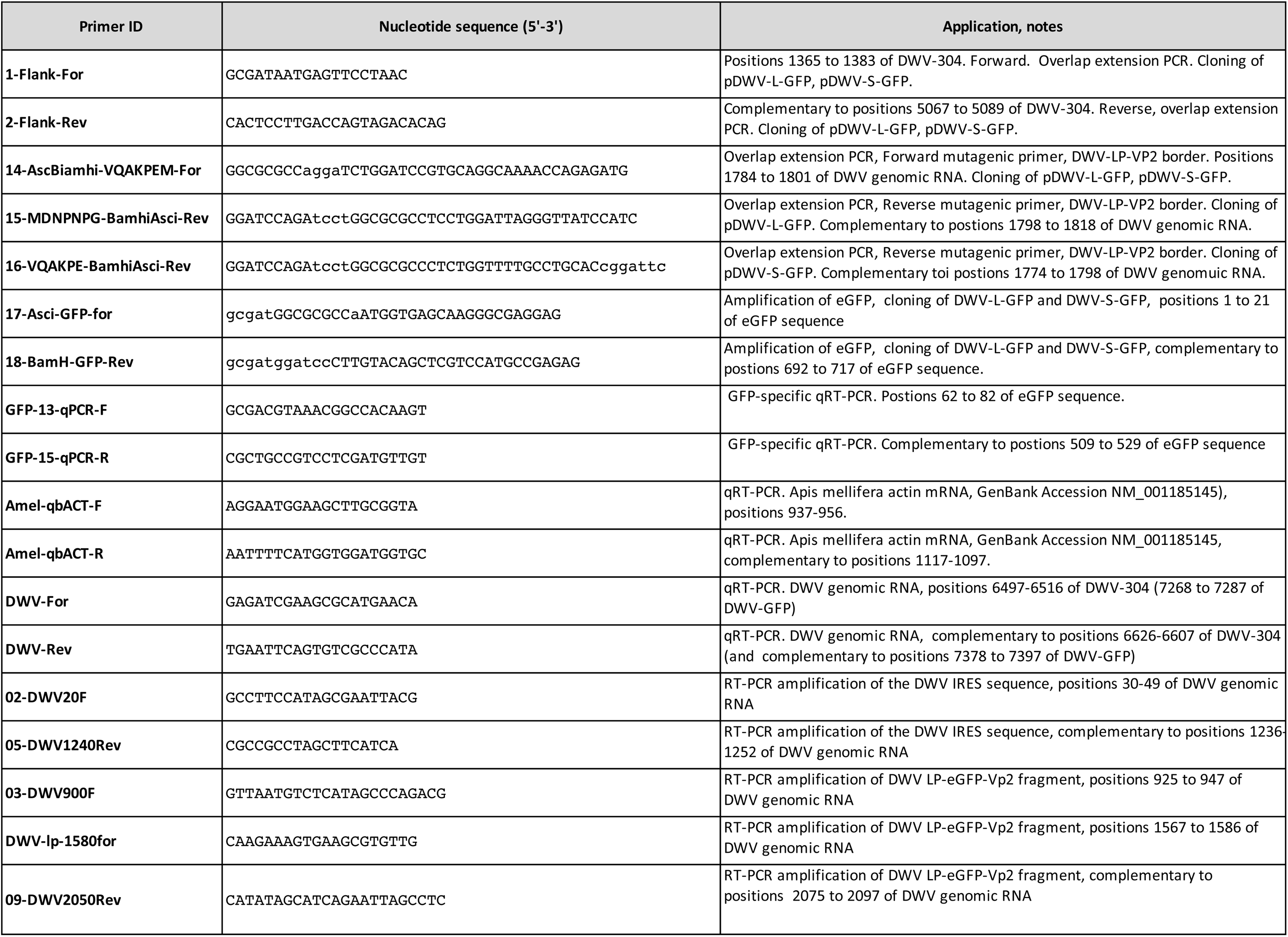
Primers and the synthetic gene used in this study. Positions in the DWV genomic RNA are acording to GenBank Accession number MG831200)

**Table S2.** Quantification of DWV and GFP RNA targets in individual pupae. Numeric values for Figure 2. ND - not determined, pupa used to prepare filtered extracts marked with *.

| Treatment, Group Number | Treatment, hours post injection (hpi). | Log 10 [DWV copies per bee] | Average Log 10 [DWV copies per bee] | SD Log 10 [DWV copies per bee] | Log 10 [GFP copies per bee] | Average Log 10 [GFP copies per bee] | SD Log 10 [GFP copies per bee] | Honey bee Actin mRNA RT-qPCR Ct values | Average Honey bee Actin mRNA RT-qPCR Ct values | SD Honey bee Actin mRNA RT-qPCR Ct values |
| --- | --- | --- | --- | --- | --- | --- | --- | --- | --- | --- |
| 1 | PBS 72 hpi | 6.489 | 5.7908 | 0.8014 | ND | ND | ND | 26.46 | 26.5943 | 0.2200 |
| 1 | PBS 72 hpi | 6.287 |  |  | ND |  |  | 26.44 |  |  |
| 1 | PBS 72 hpi | 4.480 |  |  | ND |  |  | 26.29 |  |  |
| 1 | PBS 72 hpi | 5.972 |  |  | ND |  |  | 26.64 |  |  |
| 1 | PBS 72 hpi | 4.932 |  |  | ND |  |  | 26.90 |  |  |
| 1 | PBS 72 hpi | 6.585 |  |  | ND |  |  | 26.84 |  |  |
| 2 | DWV-304 Transcript - 72 hpi | 11.094 | 11.1414 | 0.3377 | ND | ND | ND | 26.16 | 26.6093 | 0.5385 |
| 2 | DWV-304 Transcript - 72 hpi | 11.317 |  |  | ND |  |  | 26.37 |  |  |
| 2 | DWV-304 Transcript - 72 hpi | 10.510 |  |  | ND |  |  | 26.95 |  |  |
| 2 | DWV-304 Transcript - 72 hpi | 11.314 |  |  | ND |  |  | 27.49 |  |  |
| 2 | DWV-304 Transcript - 72 hpi | 11.472 |  |  | ND |  |  | 26.07 |  |  |
| 3 | DWV-L-GFP -Transcript 72 hpi* | 10.627 | 10.5402 | 0.0896 | 10.718 | 10.5132 | 0.1494 | 26.74 | 26.4996 | 0.2499 |
| 3 | DWV-L-GFP -Transcript 72 hpi | 10.417 |  |  | 10.366 |  |  | 26.15 |  |  |
| 3 | DWV-L-GFP -Transcript 72 hpi | 10.577 |  |  | 10.455 |  |  | 26.60 |  |  |
| 4 | DWV-L-GFP -Extract 72 hpi | 10.455 | 10.4603 | 0.1392 | 9.448 | 9.8704 | 0.3642 | 26.37 | 26.3124 | 0.2573 |
| 4 | DWV-L-GFP -Extract 72 hpi | 10.683 |  |  | 10.291 |  |  | 25.93 |  |  |
| 4 | DWV-L-GFP -Extract 72 hpi | 10.516 |  |  | 10.145 |  |  | 26.50 |  |  |
| 4 | DWV-L-GFP -Extract 72 hpi | 10.495 |  |  | 9.958 |  |  | 26.04 |  |  |
| 4 | DWV-L-GFP -Extract 72 hpi | 10.220 |  |  | 10.073 |  |  | 26.68 |  |  |
| 4 | DWV-L-GFP -Extract 72 hpi | 10.392 |  |  | 9.308 |  |  | 26.37 |  |  |
| 5 | DWV-S-GFP -Transcript 72 hpi | 10.815 | 10.670 | 0.1146 | 10.599 | 10.545 | 0.1293 | 26.80 | 26.276 | 0.3031 |
| 5 | DWV-S-GFP -Transcript 72 hpi | 10.495 |  |  | 10.326 |  |  | 26.14 |  |  |
| 5 | DWV-S-GFP -Transcript 72 hpi | 10.668 |  |  | 10.594 |  |  | 26.12 |  |  |
| 5 | DWV-S-GFP -Transcript 72 hpi* | 10.701 |  |  | 10.661 |  |  | 26.05 |  |  |
| 6 | DWV-S-GFP -Extract 6 hpi | 6.352 | 6.4111 | 0.0757 | 6.546 | 6.4078 | 0.1209 | 26.03 | 26.0789 | 0.2010 |
| 6 | DWV-S-GFP -Extract 6 hpi | 6.307 |  |  | 6.450 |  |  | 25.83 |  |  |
| 6 | DWV-S-GFP -Extract 6 hpi | 6.513 |  |  | 6.512 |  |  | 26.24 |  |  |
| 6 | DWV-S-GFP -Extract 6 hpi | 6.474 |  |  | 6.284 |  |  | 26.37 |  |  |
| 6 | DWV-S-GFP -Extract 6 hpi | 6.409 |  |  | 6.247 |  |  | 25.92 |  |  |
| 7 | DWV-S-GFP -Extract 24 hpi | 9.457 | 9.4820 | 0.1570 | 8.807 | 9.1899 | 0.2612 | 26.52 | 26.4162 | 0.4031 |
| 7 | DWV-S-GFP -Extract 24 hpi | 9.672 |  |  | 8.808 |  |  | 26.42 |  |  |
| 7 | DWV-S-GFP -Extract 24 hpi | 9.265 |  |  | 9.233 |  |  | 25.87 |  |  |
| 7 | DWV-S-GFP -Extract 24 hpi | 9.536 |  |  | 9.464 |  |  | 26.01 |  |  |
| 7 | DWV-S-GFP -Extract 24 hpi | 9.483 |  |  | 9.254 |  |  | 26.49 |  |  |
| 7 | DWV-S-GFP -Extract 24 hpi | 9.273 |  |  | 9.260 |  |  | 27.23 |  |  |
| 7 | DWV-S-GFP -Extract 24 hpi | 9.688 |  |  | 9.504 |  |  | 26.39 |  |  |
| 8 | DWV-S-GFP -Extract 48 hpi | 10.293 | 10.3219 | 0.1753 | 9.200 | 9.7370 | 0.2361 | 26.48 | 26.4425 | 0.2196 |
| 8 | DWV-S-GFP -Extract 48 hpi | 10.331 |  |  | 9.843 |  |  | 26.32 |  |  |
| 8 | DWV-S-GFP -Extract 48 hpi | 10.515 |  |  | 9.863 |  |  | 25.98 |  |  |
| 8 | DWV-S-GFP -Extract 48 hpi | 10.324 |  |  | 9.889 |  |  | 26.48 |  |  |
| 8 | DWV-S-GFP -Extract 48 hpi | 10.486 |  |  | 9.914 |  |  | 26.50 |  |  |
| 8 | DWV-S-GFP -Extract 48 hpi | 9.938 |  |  | 9.625 |  |  | 26.66 |  |  |
| 8 | DWV-S-GFP -Extract 48 hpi | 10.367 |  |  | 9.825 |  |  | 26.68 |  |  |
| 9 | DWV-S-GFP -Extract 72 hpi | 10.000 | 10.5222 | 0.2876 | 9.421 | 9.8058 | 0.1996 | 26.87 | 26.4025 | 0.3372 |
| 9 | DWV-S-GFP -Extract 72 hpi | 10.657 |  |  | 9.905 |  |  | 26.69 |  |  |
| 9 | DWV-S-GFP -Extract 72 hpi | 10.541 |  |  | 9.762 |  |  | 26.08 |  |  |
| 9 | DWV-S-GFP -Extract 72 hpi | 10.613 |  |  | 10.123 |  |  | 26.08 |  |  |
| 9 | DWV-S-GFP -Extract 72 hpi | 11.024 |  |  | 9.917 |  |  | 25.94 |  |  |
| 9 | DWV-S-GFP -Extract 72 hpi | 10.379 |  |  | 9.736 |  |  | 26.57 |  |  |
| 9 | DWV-S-GFP -Extract 72 hpi | 10.441 |  |  | 9.776 |  |  | 26.60 |  |  |

**Table S3.**
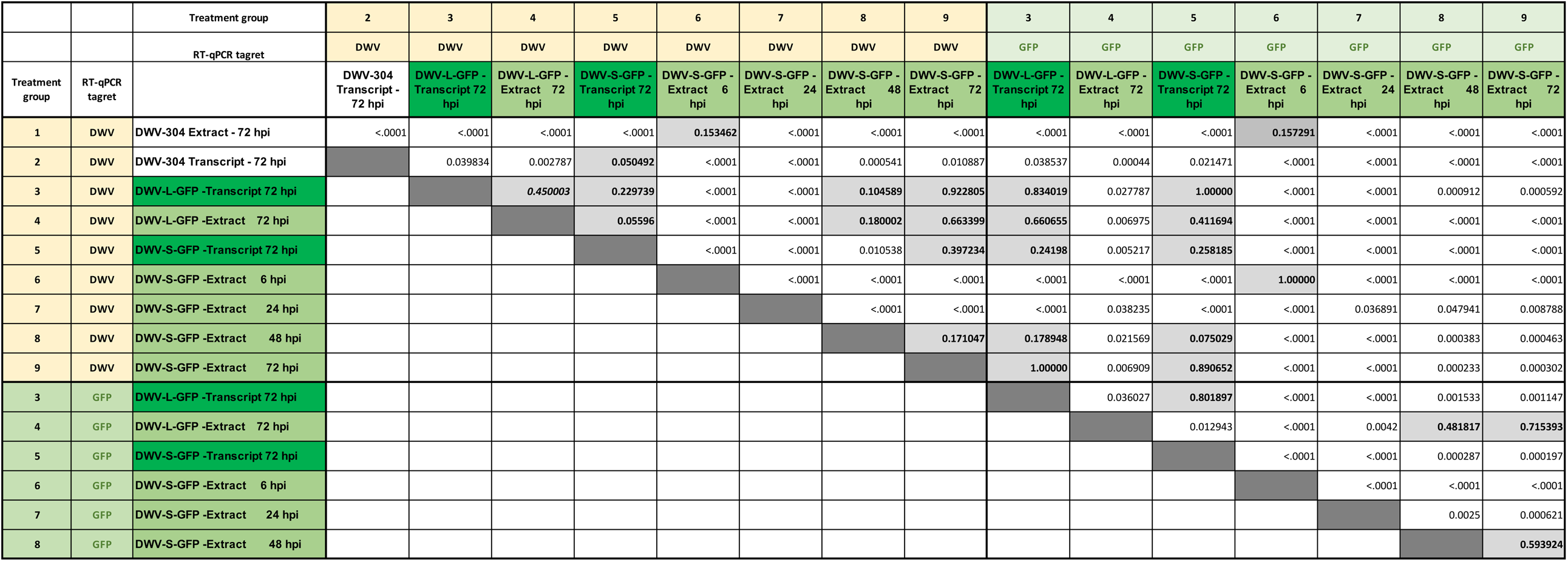
Statistical significance of the differences between DWV and GFP RNA copy numbers. P-values, ANOVA analysis. Non-significant (P>0.05%, ANOVA) are highlighted.

## Text S1. Design of pDWV-L-GPF and DWV-S-GFP

The full-length infectious cDNA clone of DWV type A, pDWV-304 (GenBank accession number MG831200) [Rybov et al., 2019] was used to design eGFP-expressing viral constructs, pDWV-L-GFP and pDWV-S-GFP (Figure 1; Text S1). Cloning included the following steps. At the first stage, two DNA fragments corresponding to the nucleotide positions 1365 – 5089 of DWV-304 with the insertion of the sequence containing restriction sites *AscI* and *BamH*I at the LP-VP2 were generated by overlap extension PCR using the plasmid pDWV-304 as a template and the primers listed in Table S1. Both fragments had *Asc*I*-Bam*HI sequences inserted upstream the sequence coding for the proposed proteolytic cleavage peptide (VQAKPEMDNPNPG) between the LP and VP2 of DWV-304. Towards the 5’ end from the *Asc*I site, the sequence coding for entire proposed proteolytic cleavage peptide at the DWV LP-VP2 interface (VQAKPEMDNPNPG) was inserted in the fragment destined for the construct pDWV-L-GFP, and the sequence coding for the proposed carboxy-terminal peptide of the DWV LP (VQAKPE) was inserted for fragment used in designing pDWV-S-GFP. As a result, the *Asc*I-*Bam*Hi sequence was linked with the sequences coding for the potential DWV LP-VP2 proteolytic cleavage peptides (Figure S1). For the pDWV-L-GFP design, the this fragment was generated by overlap extension PCR using mutagenic primers “14-AscBiamhi-VQAKPEM-For” and “15-MDNPNPG-BamhiAsci-Rev” and the flanking primers “1-Flank-For” and “2-Flank-Rev”. For the pDWV-S-GFP design, the fragment was generated using mutagenic primers “14-AscIBamhi-VQAKPEM-For” and “16-VQAKPE-BamhiAsci-Rev”, and the flanking primers “1-Flank-For” and “2-Flank-Rev”. The resulting fragments were cloned into pCRII-TOPO vector (Invitrogen) and were used to insert the *Asc*I-*Bam*HI-flanked eGFP PCR fragment generated using the primers “17-Asci-GFP-for” and “18-BamH-GFP-Rev” and pAcP(+)IE1-eGFP [Shi et al., 2007] as a template to make clones with two types of the LP-eGFP-VP2 insert. Then *Sal*I-*Sac*II section of these DWV-LP-eGFP-VP2 clones were used to replace *Sal*I-*Sac*II section of the pDWV-304 (positions 1438-4273) to produce pDWV-L-GFP and pDWV-S-GFP (Fig. 1).

## Text S2. Nucleotide sequences of the Deformed wing virus vector cDNA clones pDWV-L-GFP and pDWV-S-GFP

The clones are based on the clone DWV-304 (GenBank Accession number MG831200). GFP RNA inserts are shown in green, yellow highlight – duplicated flanking sequences derived from the DWV LP-VP2 border, blue highlight – sequences of *Asc*I and *Bam*HI restriction sites.

### >DWV-L-GFP

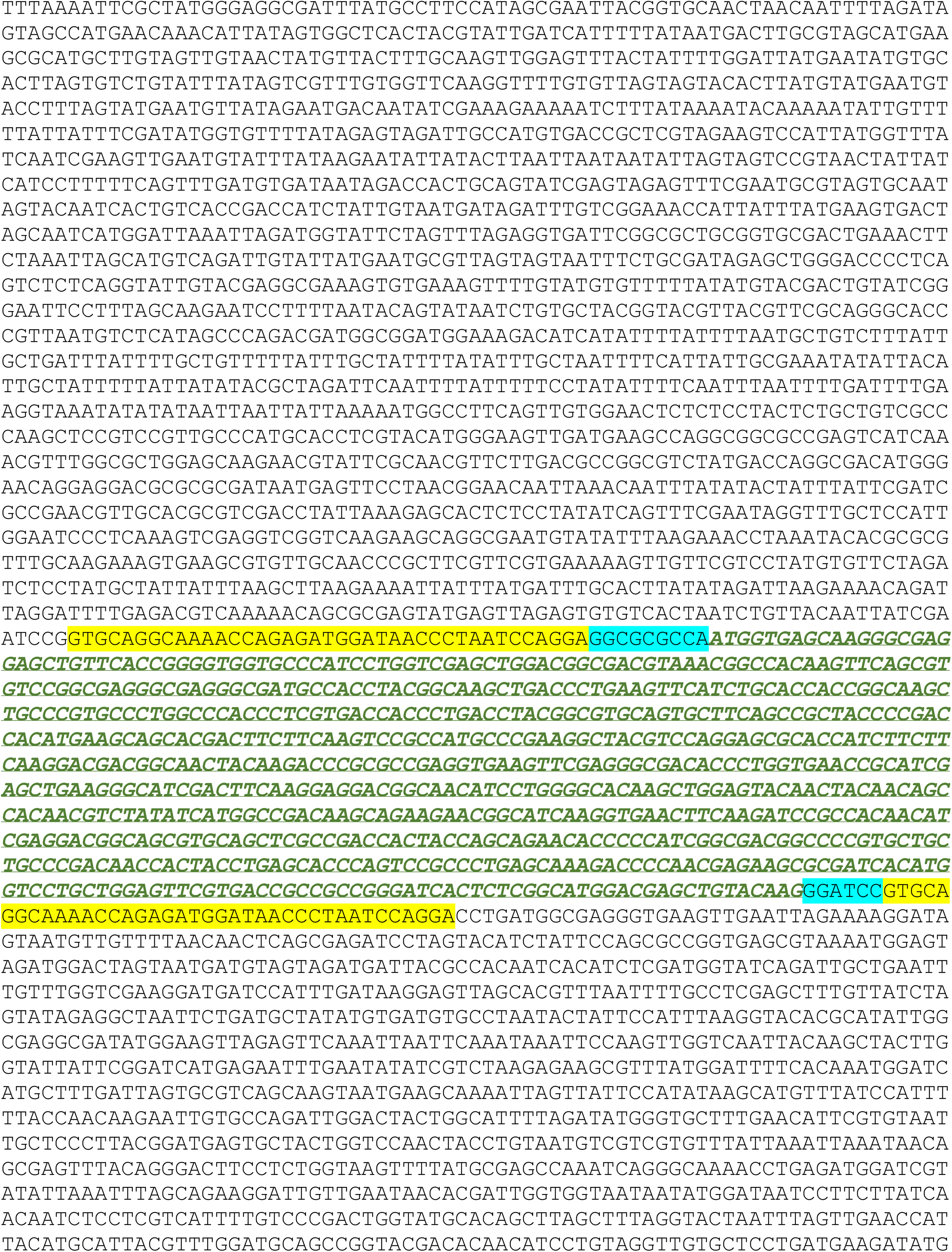
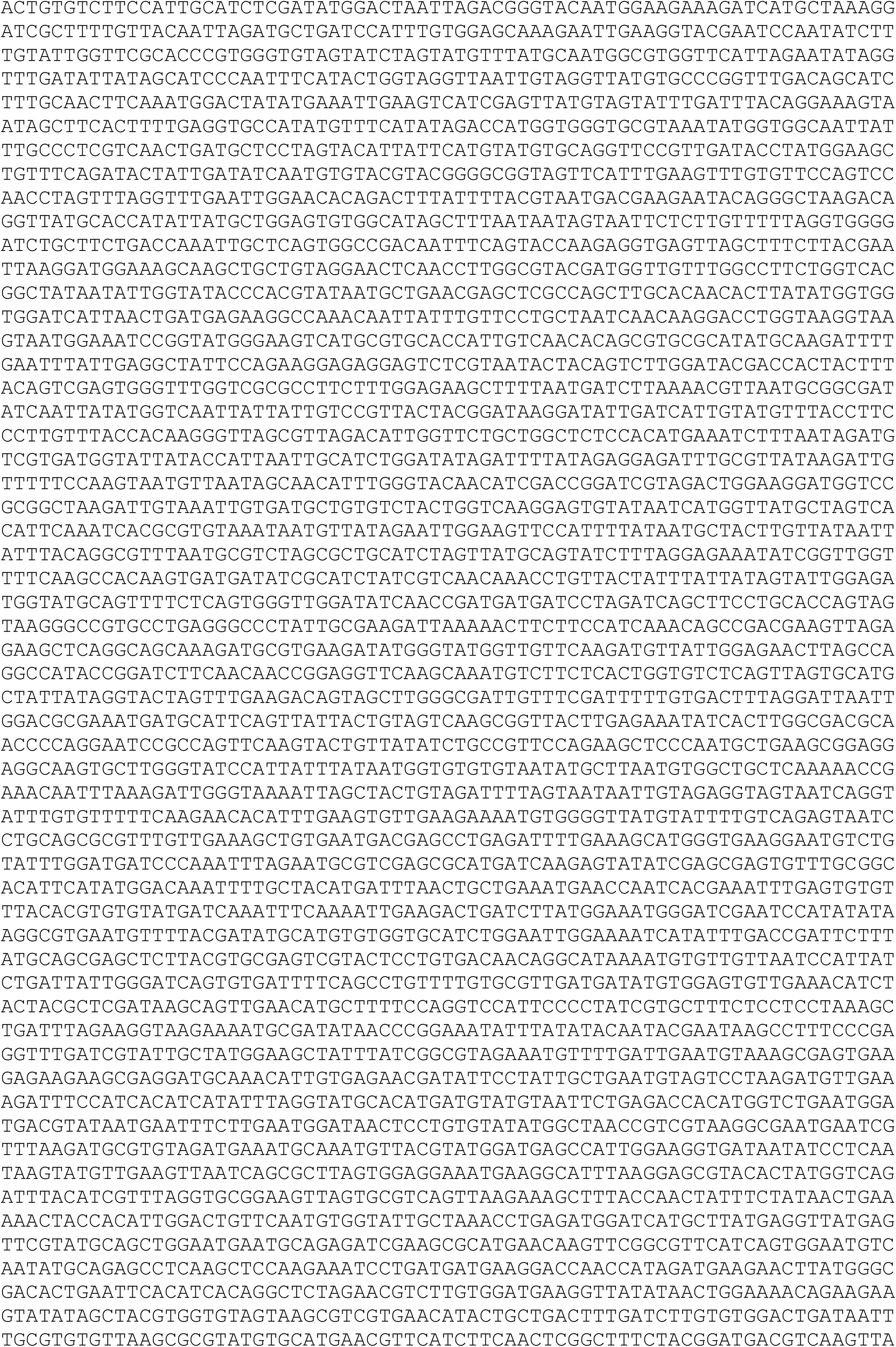
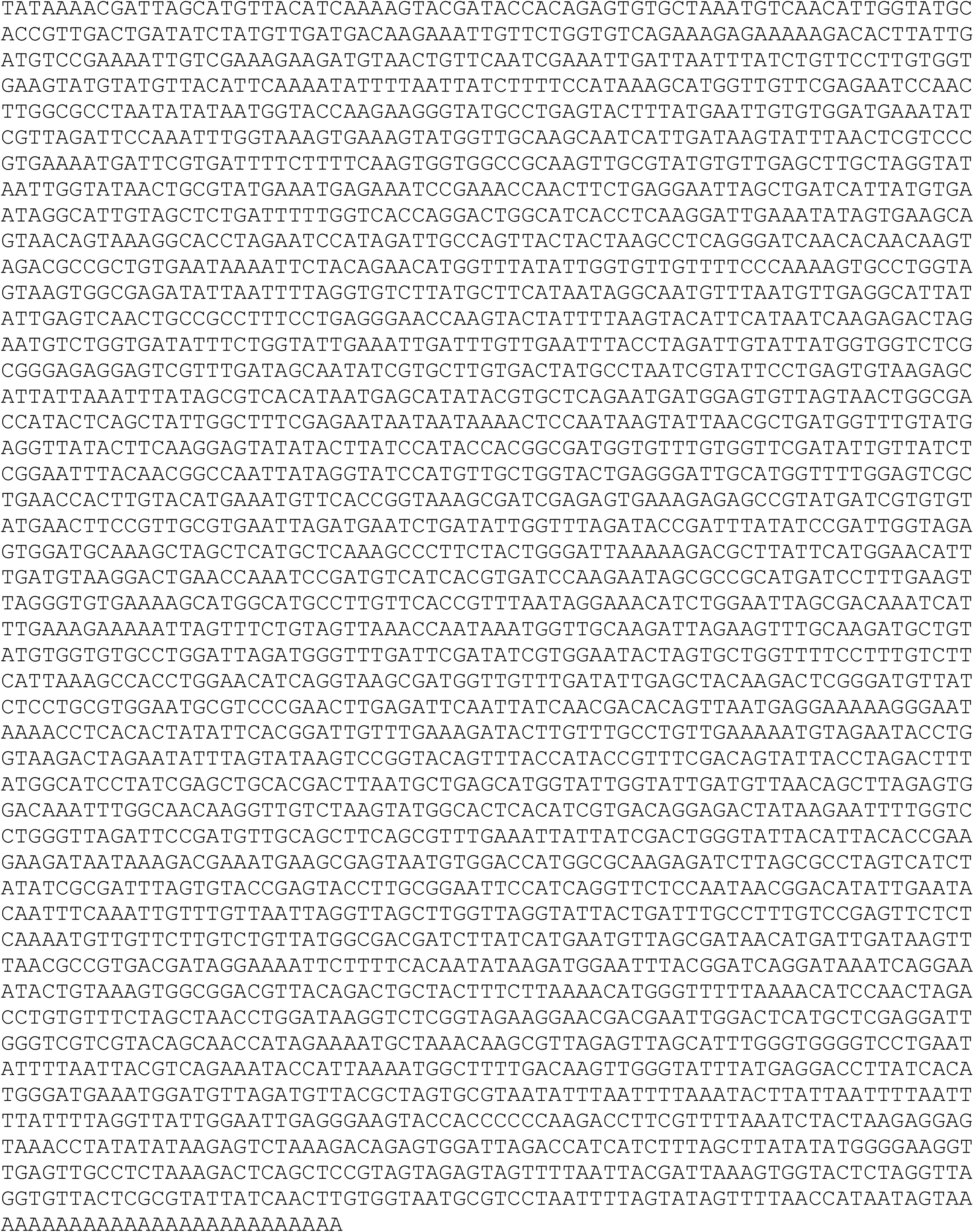

### >DWV-S-DWV

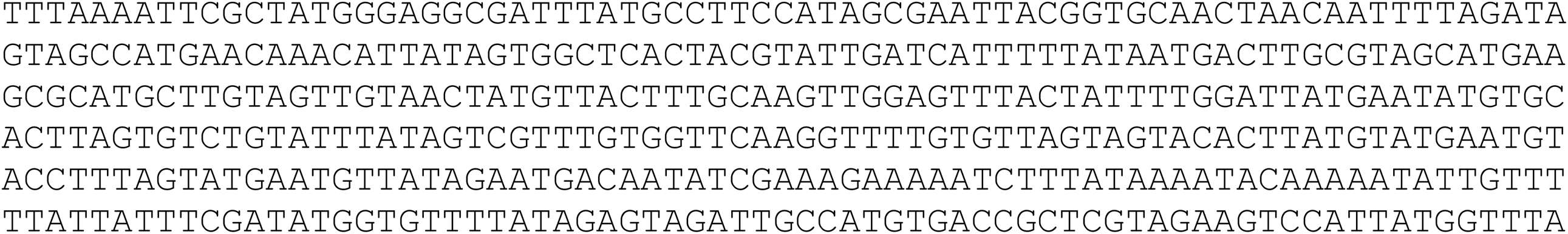
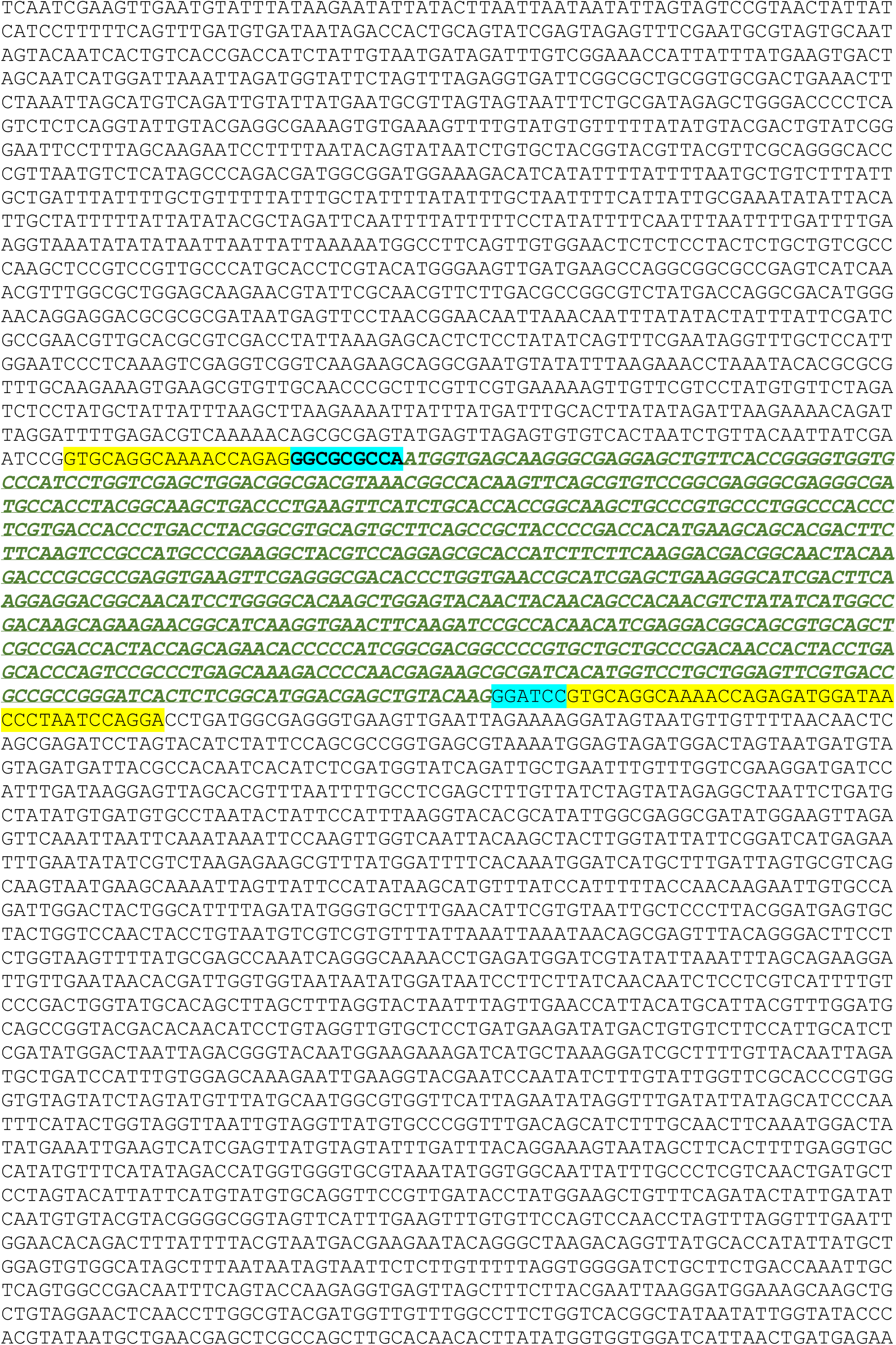
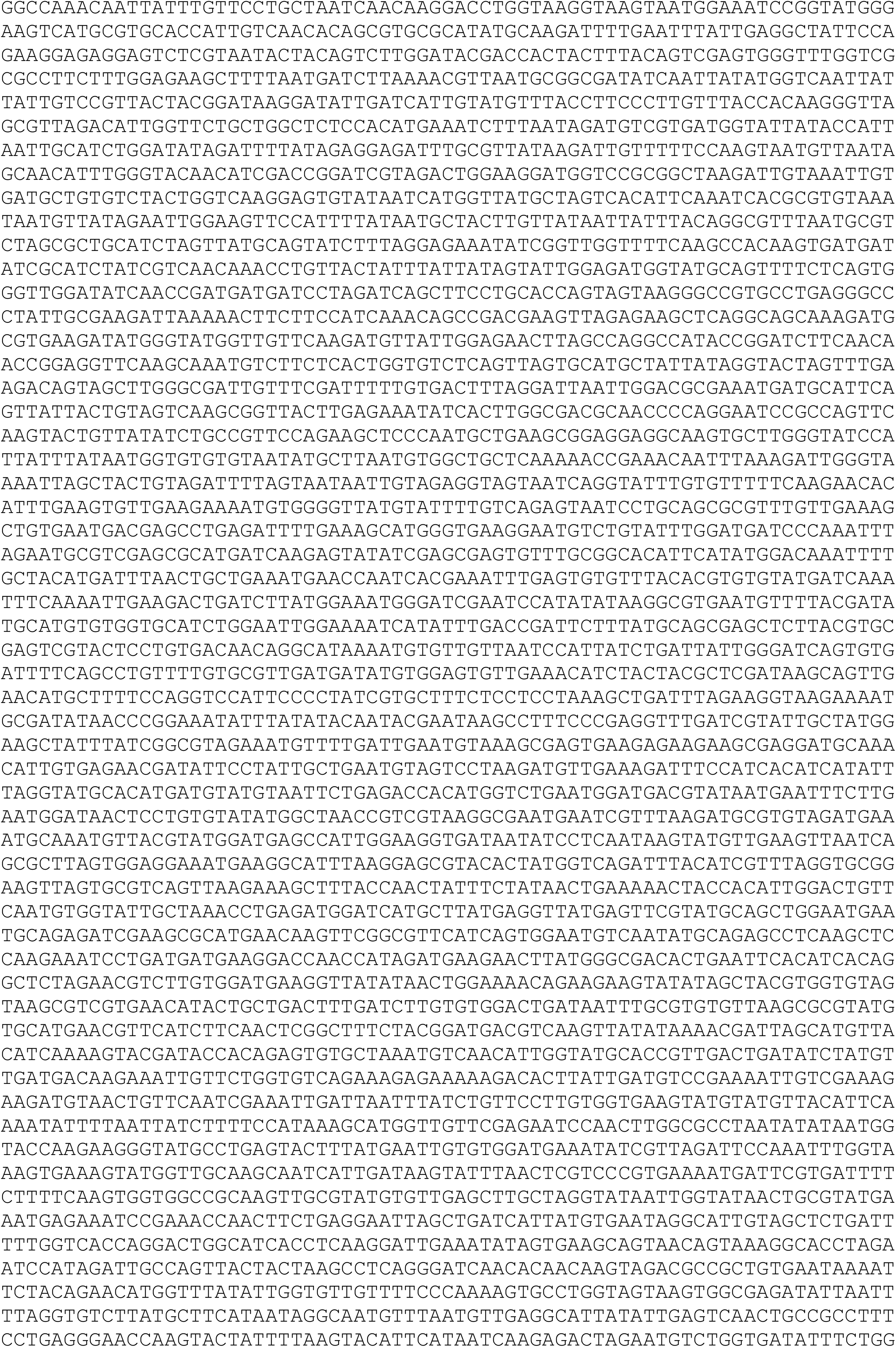
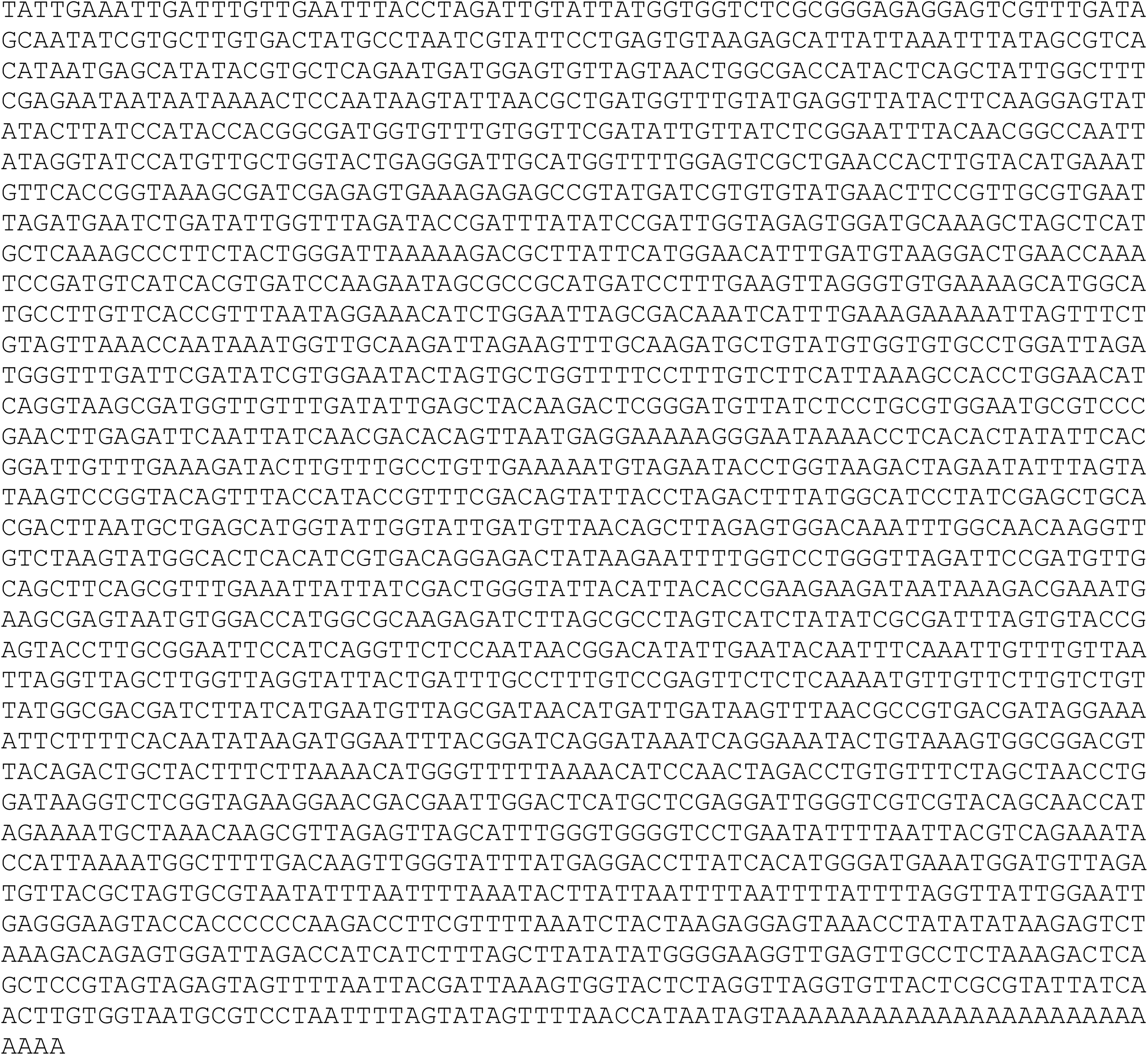

